# A platform for brain-wide functional ultrasound imaging and analysis of circuit dynamics in behaving mice

**DOI:** 10.1101/2020.04.10.035436

**Authors:** Clément Brunner, Micheline Grillet, Arnau Sans-Dublanc, Karl Farrow, Théo Lambert, Emilie Macé, Gabriel Montaldo, Alan Urban

## Abstract

Imaging of large-scale circuit dynamics is crucial to gain a better understanding of brain function, but most techniques have a limited depth of field. Here we describe vfUSI, a platform for brain-wide volumetric functional ultrasound imaging of hemodynamic activity in awake head-fixed mice. We combined high-frequency 1024-channel 2D-array transducer with advanced multiplexing and high-performance computing for real-time 3D Power Doppler imaging at high spatiotemporal resolution (220×280×175-μm^3^ voxel size, up to 6 Hz). In addition, we developed a standardized software pipeline for registration and segmentation based on the Allen Mouse Common Coordinate Framework, allowing for temporal analysis in 268 individual brain regions. We demonstrate the high sensitivity of vfUSI in multiple experimental situations where stimulus-evoked activity can be recorded using a minimal number of trials. We also mapped neural circuits *in vivo* across the whole brain during optogenetic activation of specific cell-types. Moreover, we revealed the sequential activation of sensory-motor regions during a grasping water droplet task. vfUSI will become a key neuroimaging technology because it combines ease of use, reliability, and affordability.

## INTRODUCTION

Neuroscientists are continuously refining the modular description of the brain into anatomically and functionally distinct regions. Still, recent studies have reported that even simple tasks require interactions between many cortical (Ferezou et al., 2007) and subcortical areas (Steinmetz et al., 2019). How neuronal activity is coordinated between cortical and subcortical regions to produce behavior is still poorly understood in part due to the lack of technologies to monitor activity across the whole brain, especially in mice that are extensively used for neuroscience research. Optical methods are widely used for imaging both blood flow and electrical responses of the brain in awake mice. Nevertheless, existing imaging strategies remain challenging to apply in awake conditions or lack the penetration and resolution necessary for precise identification of neuronal circuits spanning both cortical and subcortical brain regions (Urban et al., 2017). Moreover, recently developed multielectrode array allows recordings of both local field and action potentials from hundreds of neurons distributed across several brain regions but remains invasive and with a limited brain coverage (Jun et al., 2017).

To date, fMRI is the only available methodology for probing large scale functional circuits in the whole mouse brain in awake condition. However, performing mouse fMRI needs expensive ultra-high-field scanners and cryogenic coils (Uludağ and Blinder, 2018) and has a limited spatiotemporal resolution. Recent studies have reported fMRI in resting condition (Yoshida et al., 2016), during somatosensory (Desjardins et al., 2019; Harris et al., 2015) as well as during an olfactory go/no-go task (Han et al., 2019) in an awake mouse. fMRI was combined with optogenetics to investigate the functional connectivity of precise neuronal circuits across the entire intact brain at low temporal resolution (Desai et al., 2011). The fact that most available results are, however, acquired under anesthesia, which significantly limits the scope of their application and behavioral relevance (Baltes et al., 2011; Komaki et al., 2016; Shim et al., 2018).

Functional ultrasound imaging (fUSI) is a method based on high-framerate plane-waves imaging to measure changes in cerebral blood volume (CBV) in response to neuronal activity (Mace et al., 2013; Mace E. et al., 2011; Montaldo et al., 2009; Urban et al., 2014a, 2017). fUSI has a small voxel size (~100×300×100 μm^3^), an excellent temporal resolution (10 Hz framerate), a high depth of field (up to 1.5 cm), does not require contrast agents and is adaptable to freely-moving animals (Sieu et al., 2015; Tiran et al., 2017; Urban et al., 2015). Nevertheless, fUSI was initially restricted to crosssectional images. Visualizing the whole brain requires stepping the probe across multiple positions, as it was recently shown to identify brain regions involved in the optokinetic reflex in awake mice (Macé et al., 2018). This strategy, however, results in long acquisition time and repetitive presentation of the stimulus at each position, which may hinder animal physiology and trigger habituation (Miller et al., 2018).

The initial proof of concept study showed that 4D fUSI (Rabut et al., 2019) could potentially overcome the current limitations of 2D fUSI in terms of acquisition speed. However, this result was achieved at the cost of decreasing the spatiotemporal resolution and using experimental conditions limiting the usefulness of 3D imaging. Our main objective was to develop a volumetric functional ultrasound imaging (vfUSI) platform with specifications that could meet 2D-fUSI standard hence overpassing current issues of 4D fUSI.

First, we improved the imaging framerate (from 0.3 to up to 6 Hz), voxel size (from 320×320×180 to 220×280×175 μm^3^), and sensitivity by increasing ultrasound transducer frequency (from 8 to 15 MHz, 300μm pitch) and adapting both ultrasound electronics and compound plane-waves ultrasound sequence. Second, we resolved experimental limitations due to general anesthesia use to limit brain motion artifacts during imaging. Indeed, medetomidine/ketamine may reduce fUSI signal relevance because it impairs neurovascular coupling (Masamoto and Kanno, 2012). The solution was to implement a reliable cranial window surgical procedure based on our 2D fUSI knowledge (Brunner et al., 2017; Macé et al., 2018) that is suitable for chronic (up to five months) brain-wide vfUSI in awake behaving mice. This strategy was efficient in decreasing the number of animals to a minimum and allowed repeated measurement, hence improving data quality and relevance by reducing inter-subject variability in brain function (Seghier and Price, 2018). Third, we wanted to ensure the widespread availability of this technology to the neuroscience community by selecting only commercially available instead of proprietary hardware components like in 4D fUSI (Provost et al., 2014). We aimed at validating vfUSI in behaving mice. In fact, mice are leading model organisms for neuroscience research instigated by the availability of a much larger genetic toolbox for manipulating and recording neurons than in rats (Luo et al., 2008), providing more direct translational opportunities regarding brain functional comparison between human and rodent research. Nevertheless, developing vfUSI in behaving mice created additional difficulties as compared to anesthetized rats due to the small size of the brain but also because of motion artifacts (Desjardins et al., 2019). Moreover, we have devoted particular attention to standardizing vfUSI data acquisition and analysis mimicking routinely used procedures in fMRI (Lindquist, 2008). For this purpose, we developed a dedicated software pipeline for quick, easy, and reliable data registration based on the Allen Mouse Common Coordinate Framework (http://atlas.brain-map.org/), providing hemodynamic measurements across 268 individual brain regions.

Several experiments were performed in awake mice to validate our innovative vfUSI platform. We assessed the resolution and specificity of vfUSI by stimulating single whiskers and imaging the somatotopic organization of the barrel field cortex. Then, brain-wide maps of the visual system were successfully established with just a few trials averaging. Furthermore, we demonstrated that vfUSI could be efficiently combined with optogenetic stimulation. Here, three distinct cell types in the superior colliculus were identified to reveal the differences in projection patterns and activity across several synapses. As the last step, we recorded brain activity during an active sensory-motor task and highlighted the ability of vfUSI to capture distributed neuronal information with complex dynamics at large scale.

## RESULTS

### vfUSI hardware and software components for real-time acquisition and processing

We developed a 2D-array transducer with 1024 elements with an active head surface of 9.6×9.6 mm^2^ (Vermon, France) covering a large part of the mouse brain. Importantly, the design of the piezoelectric ultrasound array transducer is constrained by current manufacturing and wiring processes, limiting the number of channels to ~1000 and inter-element spacing to 300 μm in both azimuth (x) and elevation (y) directions. Considering that the reflectivity of red blood cells highly increases with frequency (Yu and Cloutier, 2007), we chose a 15-MHz (8-22 MHz bandwidth) central excitation frequency resulting in an optimal signal-to-noise ratio (SNR) and a good penetration of the ultrasound beam (up to 1.5 cm depth). The 1024 channels are organized in 4 rectangular sectors composed of 256 (8×32) elements individually wired to a 1024 to 256 (4:1) multiplexer controlled by 256 emission/reception channels ultrasound electronics (Verasonics, USA). The signal from backscattered echoes was sampled at 60 MHz, digitized at 14 bits, and transferred to a high-performance computing workstation for real-time GP-GPU processing (**Figure 1A**).

**Figure 1.**
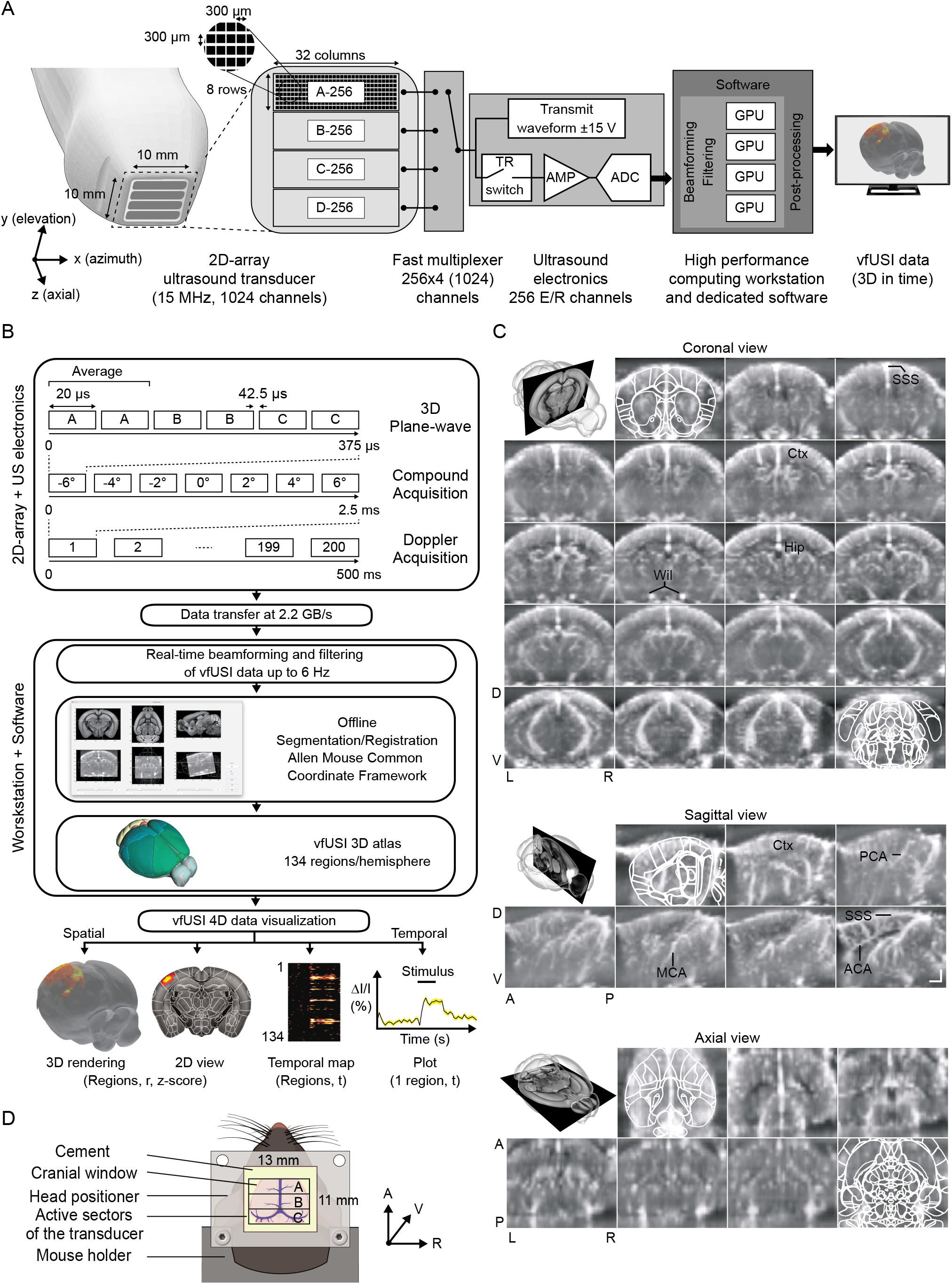
vfUSI hardware and software components for real-time acquisition and processing. **(A**) Schematic representation of the 15 MHz 2D-array transducer with a 9.6×9.6 mm^2^ active head composed of 1024 (256×4, A to D) elements spaced 300 μm apart in both azimuth (x) and elevation (y) directions. The 2D-array transducer is wired to a 4:1 fast multiplexer (1024 to 256) controlled by 256 channels ultrasound electronics. Backscattered echoes were sampled, digitalized and transferred to a high-performance computing workstation with 4 GPUs for real-time beamforming, filtering and processing of the vfUSI data. **(B**) Each vfUSI image was generated from backscattered echoes (20 μs) averaged 2 times to improve the signal to noise ratio. 200 compound images (7 angles from −6° to 6°) were used to generate a Doppler image in 500 ms. 200 ms were necessary to beamform and filter the vfUSI data in real-time with a bandwidth of ~2.2 GB/s corresponding to an imaging frequency varying from 1.4 Hz up to 6 Hz. Segmentation and registration of the 3D vfUSI data were performed in 134 brain regions/hemisphere using a dedicated software based on the Allen Mouse Common Coordinate Framework. **(C**) Various cross-sectional views extracted from the vfUSI data. Top panel: coronal view (anterior to posterior). Middle panel: sagittal view (lateral to medial). Bottom panel: axial view (dorsal to ventral). An example of registration on the Allen Mouse Brain Atlas was outlined in white. Some typical regions, such as the cortex (Ctx), the hippocampus (Hip), and main vessels such as Anterior, Middle and Posterior Cerebral Arteries (A/M/PCA), Circle of Willis (Wil) and Superior Sagittal Sinus (SSS) are highlighted in black. **(D)** A rectangular stainless-steel head positioner with a large imaging window (13×11 mm^2^) was cemented on the mouse’ skull bone for vfUSI in awake head-fixed conditions. After applying agarose and ultrasound gel to improve the acoustic coupling, the 2D-array transducer was placed over the brain. The number and position of active sectors (A-C) were adapted to each experimental paradigm (**Table S1**). A, anterior; P, posterior; D, dorsal; V, ventral; L, left; R, right. Scale bars, 1 mm.

The vfUSI ultrasound sequence was adapted from the previously developed 2D fUSI sequence (Montaldo et al., 2009). Briefly, 3D compound plane-waves are generated by sequential emission and reception of a subset of sectors from the 2D-array. Each sector is activated two times to average the reception and increase the SNR (**Figure 1B**). Using these parameters, we obtained a voxel size of 220×280×175 μm^3^ along the *x-y-z* axes (**Figure S1A**; **STAR Methods**). The vfUSI ultrasound sequence has a ~2.2 GB/s bandwidth that is further processed online and compressed to reduce the file size of each 3D image to ~1 MB allowing data storage without any restriction on the duration of the experiment. Hence, a critical part of the ultrasound scanner development relied on the implementation of dedicated software for real-time processing of the vfUSI data. For this purpose, we combined 4 GPUs (Nvidia Geforce RTX2080Ti, USA) with massively parallel computation capabilities required for online beamforming and filtering (**Figure 1B**; **STAR Methods**). This architecture offers an efficient duty cycle of 71% (500 ms acquisition, 200 ms dead time) for recording vfUSI movies at a frame rate of ~1.4 Hz. The hemodynamic signal (Δl/l, referred to as ‘‘hemodynamic response or activity”, see **STAR Methods**), is proportional to CBV (Mace et al., 2013) and was further extracted in each voxel. Importantly, the vfUSI scanner offers excellent flexibility between sensitivity, framerate, and field of view by adjusting the number of angles, compound images, and transducer aperture. As demonstrated below, when necessary, the temporal resolution of vfUSI can be increased with a limited effect on the sensitivity (data not shown) by performing data acquisition to up to 6-Hz frame rate (100% duty cycle) using only 3 compound angles and 80 compound images.

### Animal surgery and analysis pipeline for brain-wide activity mapping

We designed an implantable stainless-steel head positioner with a large rectangular opening (13×11 mm^2^) that was cemented on the skull bone after sterilization. Consecutively, a cranial window (~8×10 mm^2^) was performed to expose the entire brain except for the olfactory bulb and the back of the cerebellum (**Figure 1D**). The dura mater was kept intact and protected with a silicone cap reducing bone regrowth (Spitler and Gothard, 2008). Before imaging sessions, the cap was removed, and the dura was covered with 2.5% agarose to dampen brain motions. This protocol enabled for brainwide vfUSI in awake head-fixed mice in chronic conditions for up to 5 months with minimal brain tissue inflammation, as previously observed in rats (Urban et al., 2014a). The 2D-array transducer was then placed ~2 mm from the brain surface, and additional ultrasound gel was applied to improve acoustic coupling. The number and positions of active sectors were adapted to each experimental paradigm (**Table 1**; **Figure S1C**; **STAR Methods**). We proceeded with the habituation and training of the animal to the head-fixation and imaging setup. Habituation reduces animal stress and is essential for having optimal and reproducible physiological conditions across animals and across imaging sessions. Habituation also reduced motion artifacts, so that rejected frames never exceeded 10% of the total number of frames during vfUSI. We developed dedicated software for 4D (3D over time) analysis, allowing quick image registration and segmentation based on the Allen Mouse Common Coordinate Framework (**Figure 1B**; Oh et al., 2014). The atlas was adapted into a simplified vfUSI atlas containing 134 brain regions per hemisphere after removal of non-vascular structures (i.e., ventricles, fiber tracts) and aggregation of small structures not resolved by our method (**Table S1; STAR Methods**). After the acquisition, 3D images were resampled to a 50×50×50 μm^3^ voxel size by 3D cubic interpolation and then registered by affine transformation to the vfUSI atlas using a dedicated graphical user interface (**Figure 1B**, central panel). The processed vfUSI data can be displayed in various ways, such as i) correlation maps of hemodynamic activity in cross-sections, ii) 3D heat-maps, iii) pattern of activity among all regions in each hemisphere and iv) temporal plots averaged across several voxels (**Figure 1B**, bottom panel). Codes and examples are available in **Supplementary Materials**. A qualitative analysis of cross-sectional Power Doppler images confirmed that the brain microvasculature is distinctively observable in several regions, including the cortex (Ctx) and the hippocampus (Hip). Also, large blood vessels could be visualized, such as the superior sagittal sinus (SSS), the anterior/middle/posterior cerebral arteries (A/M/P-CA), and the circle of Willis (Wil; **Figure 1C**; Xiong et al., 2017).

**Table 1 -.**
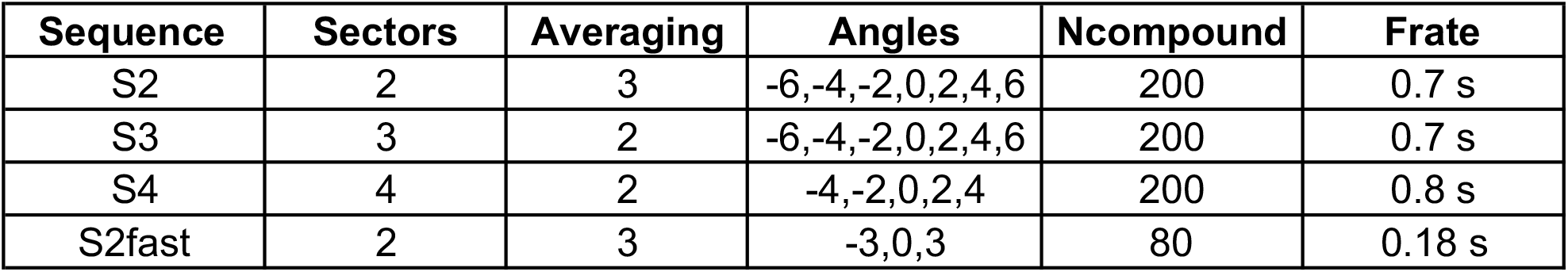
Sequences used to perform vfUSI in awake head-fixed mice and rats. Parameters are defined in **STAR Methods**.

| **Sequence** | **Sectors** | **Averaging** | **Angles** | **Ncompound** | **Frate** |
| --- | --- | --- | --- | --- | --- |
| S2 | 2 | 3 | -6,-4,-2,0,2,4,6 | 200 | 0.7 s |
| S3 | 3 | 2 | -6,-4,-2,0,2,4,6 | 200 | 0.7 s |
| S4 | 4 | 2 | -4,-2,0,2,4 | 200 | 0.8 s |
| S2fast | 2 | 3 | -3,0,3 | 80 | 0.18 s |

### Resolution, sensitivity, and specificity during whisker stimulation

vfUSI relies on a 2D-array transducer whose sensitivity, SNR and aperture strongly differ from the 1-D linear array transducer used in previous fUSI studies (Mace E. et al., 2011; Urban et al., 2015). Therefore, our foremost objective was to perform a qualitative and quantitative analysis of the *in vivo* vfUSI capabilities in awake mice. For this purpose, a sensory-evoked stimulation mimicking naturalistic deflections of multiple whiskers was used (5-Hz sinusoidal deflection, 20° amplitude, 6 s duration; **Figure 2A**). Here, we observed a statistically significant (**STAR Methods**) increase of the hemodynamic signal in the contralateral primary somatosensory barrel field cortex (SSp-bf) by averaging only five trials (**Figure 2B, left panel; Video S1**). This experiment lasted less than 3 min, potentially minimizing the effects of habituation. Nevertheless, averaging of 50 trials were necessary to detect evoked responses in the ventral posterior medial (VPM) nucleus of the thalamus because of their small amplitude (<2%) as compared to baseline hemodynamic variations. In this case, we also noticed a substantial increase of the activated volume in the cortex (~200 %) with spreading of the activation beyond the SSp-bf in both visual and auditory cortices (SSp-n, SSs, SSp-un, VISal, VISrl, VISp, AUDd, AUDpo; **Figure 2B right panel**; **Video S1**) suggesting a direct proportionality of volumetric spatial neurovascular coupling in the cerebral cortex. Temporal plots of the hemodynamic responses at the core of the activated regions showed a quick increase of the signal during whisker stimulation that reached a plateau after 1.5 s in both the cortex (5.8±0.7%) and the thalamus (1.7±0.5%; **Figure 2E**) before slowly returning to baseline. Note that the level of activation in both regions is in agreement with those already observed in previous fUSI studies in rodents (Brunner et al., 2018; Urban et al., 2015). We used the SSp-bf organization as a model to study the functional resolution and sensitivity of vfUSI to sequential stimulation of 3 individual whiskers (B2, C4, D2). We showed that stimulation of each single whisker activates non-overlapping specific regions of the barrel cortex (**Figure 2C**; **Video S1**) even if 50 repetitions were necessary to detect such small increases of CBV signal (below 2%; **Figure 2F**). We observed that the activation map of each whisker is spatially very confined (~650×600×500 μm^3^) (**Figure 2C**) but was slightly larger than the size of a single barrel (Petersen, 2007) and potentially due to a spread of the hemodynamic signal (Devor et al., 2005). In one mouse, cytochrome oxidase staining was performed on the flattened cortex to label the cortical layer IV barrels over which activated voxels were overlaid (**STAR Methods**). We confirmed that the relative position of each activated region fitted quite well with the mouse somatotopic whisker map (**Figure 2D**).

**Figure 2.**
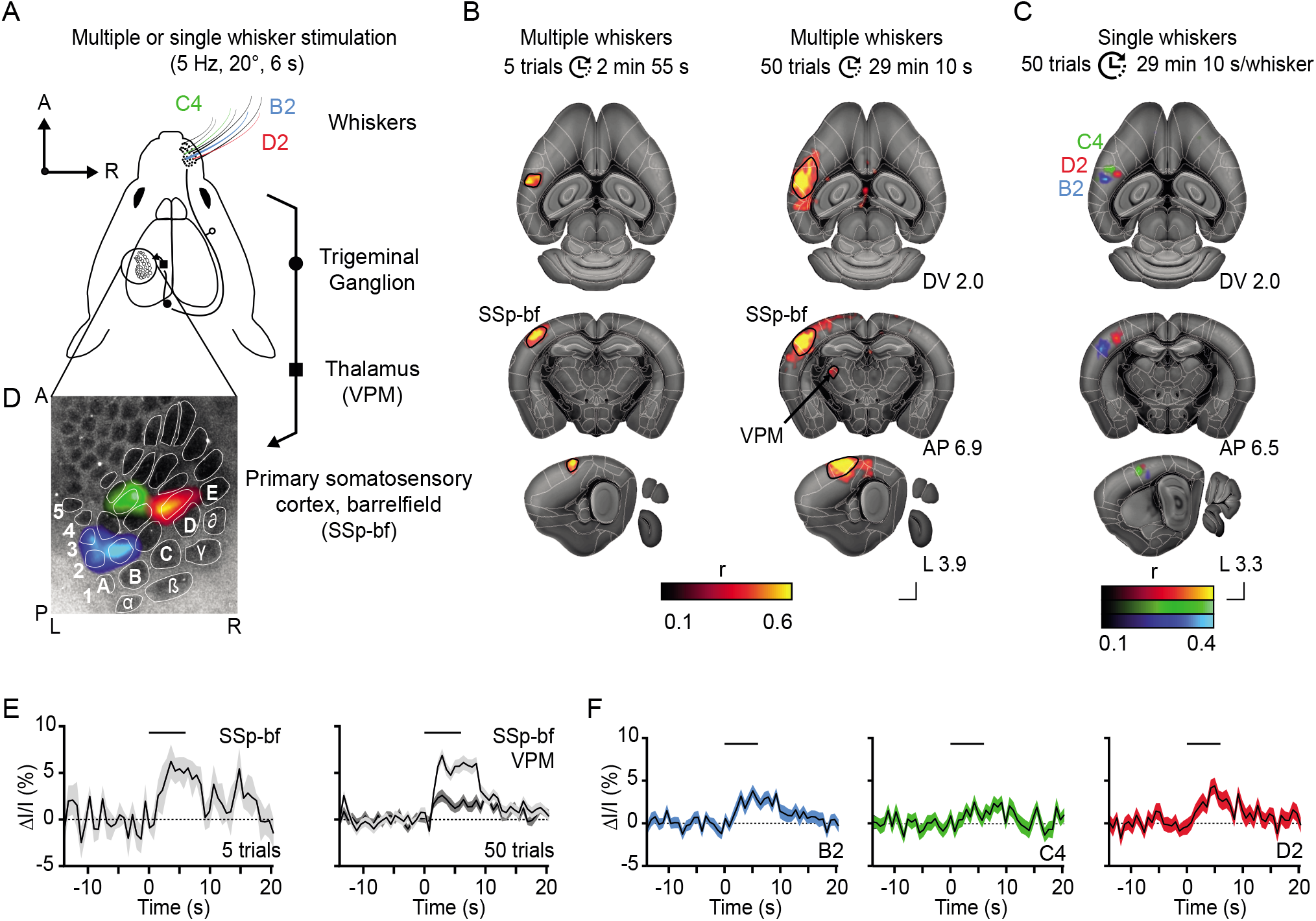
vfUSI spatial sensitivity and specificity during whisker stimulation. **(A)** Schematic of the mouse whisker to barrel pathway. Multiple or single whiskers (B2, C4, D2) were stimulated to evoke sensory hemodynamic responses. **(B)** Example slices from the 3D correlation map presented in different views (axial, coronal, and sagittal) showing activated voxels during multiple whisker stimulation (average of 5 and 50 trials) superimposed on the anatomical atlas (greyscale). We observed a statistically significant increase of activity in the contralateral SSp-bf and the VPM nuclei of the thalamus. Significant activation corresponds to r values > 0.2. **(C)** Example slices from the 3D correlation map presented in different views (axial, coronal, and sagittal) showing activated voxels during single whisker stimulation (B2, blue; C4, green and D2, red) superimposed on the anatomical atlas. Non-overlapping hemodynamic activities corresponding to each whisker were observed in the SSp-bf. **(D)** Cytochrome oxidase staining of the cortical barrels was overlaid on the vfUSI signal. **(E)** Average hemodynamic response curves extracted from activated voxels (black dotted areas in **(B)**) in response to multiple whisker stimulation in the SSp-bf and VPM for 5 (left plot) and 50 trials (right plot). Black thick line, stimulus. Grey highlighted traces, standard error of the mean. **(F)** Average hemodynamic response curves extracted from activated voxels (blue, green, and red respectively in **(C)**) in response to stimulation of individual whisker. Black thick line, stimulus. Color highlighted traces, standard error of the mean. A, Anterior; P, posterior; D, dorsal; V, ventral; L, left; R, right. Scale bars, 1 mm.

As a proof of concept, we applied vfUSI in awake head-fixed rats and performed the same experiment, but using all sectors of the 2D-array transducer (**Table 2**) due to the larger brain size. Here again, we successfully imaged evoked hemodynamic responses during both multiple and single whisker stimulations and obtained similar results to those in mice (**Figure S4**).

**Table 2.** 2D-array ultrasound transducer sequence and orientation

| Animal model | Experiment | Figure | Sequence | 2D-transducer array orientation |
| --- | --- | --- | --- | --- |
| Mouse | Doppler image | Figure 1D top | S3 | Coronal |
|  | Doppler image | Figure 1D middle | S2 | Sagittal |
|  | Whisker stimulation | Figure 2 | S2 | Coronal |
|  | Visual stimulation | Figure 3 | S2 | Coronal |
|  | Optogenetic stimulation | Figure 4 | S2 | Sagittal |
|  | Water droplet-reaching task | Figure 5 | S2fast | Coronal |
|  | RSFC | Figure S3 | S3 | Coronal |
| Rat | Whisker | Figure S2 | S4 | Coronal |

### Brain-wide vfUSI of visual-evoked activity

The next set of experiments used visual stimulation to demonstrate the efficiency of vfUSI in detecting whole-brain activations in awake head-fixed mice. First, a full-field vertical grating drifting in the naso-temporal direction to the right eye was presented during 16 s (**Figure 3A; left panel**). Ten repetitions were sufficient to obtain a robust 3D activation map (**Figure 3A; right panel** and **Video S1**). Analysis of the visually evoked responses in the contralateral hemisphere revealed that both cortical and subcortical regions were significantly activated during the stimulus including brain areas involved in visual processing such as the primary visual area of the cortex (VIS), the dorsal part of the lateral geniculate complex (LGd), the lateral posterior (LP) nucleus of the thalamus and the superior layer of the superior colliculus (SCs; **Figure 3B**). We also detected activity in regions beyond the main visual pathways, such as the inferior colliculus (IC) and the postsubiculum (POST). Averaging the hemodynamic responses in each region showed sustained increases of the CBV signal during the stimulus with an activity overshoot at the beginning and end of visual stimulation that may correspond to both ‘on’ and ‘off’ responses, respectively (**Figure 3C**; Wang et al., 2010). It should be noted that these experiments were performed in ~6 min, which is 10× faster than the time necessary to obtain similar results using the mechanical translation of a linear ultrasound transducer across the brain (Macé et al., 2018). The main difference is that we used monocular and nasotemporal visual motion stimulation, which does not elicit reflex eye movements to map only visual responses, and not oculomotor signals (Macé et al., 2018).

In the second set of visual experiments, 50 presentations of a vertical bar drifting in a horizontal direction were averaged in a total duration of 12 min 30 s per experiment (**Figure 3D**). In this condition, we were able to map the retinotopic organization of both the visual cortex and the superior colliculus. As already reported for both regions, we confirmed that the zero vertical meridian was represented rostrally, and the most peripheral part of the contralateral visual field, caudally (Drager and Hubel, 1975, 1976).

**Figure 3.**
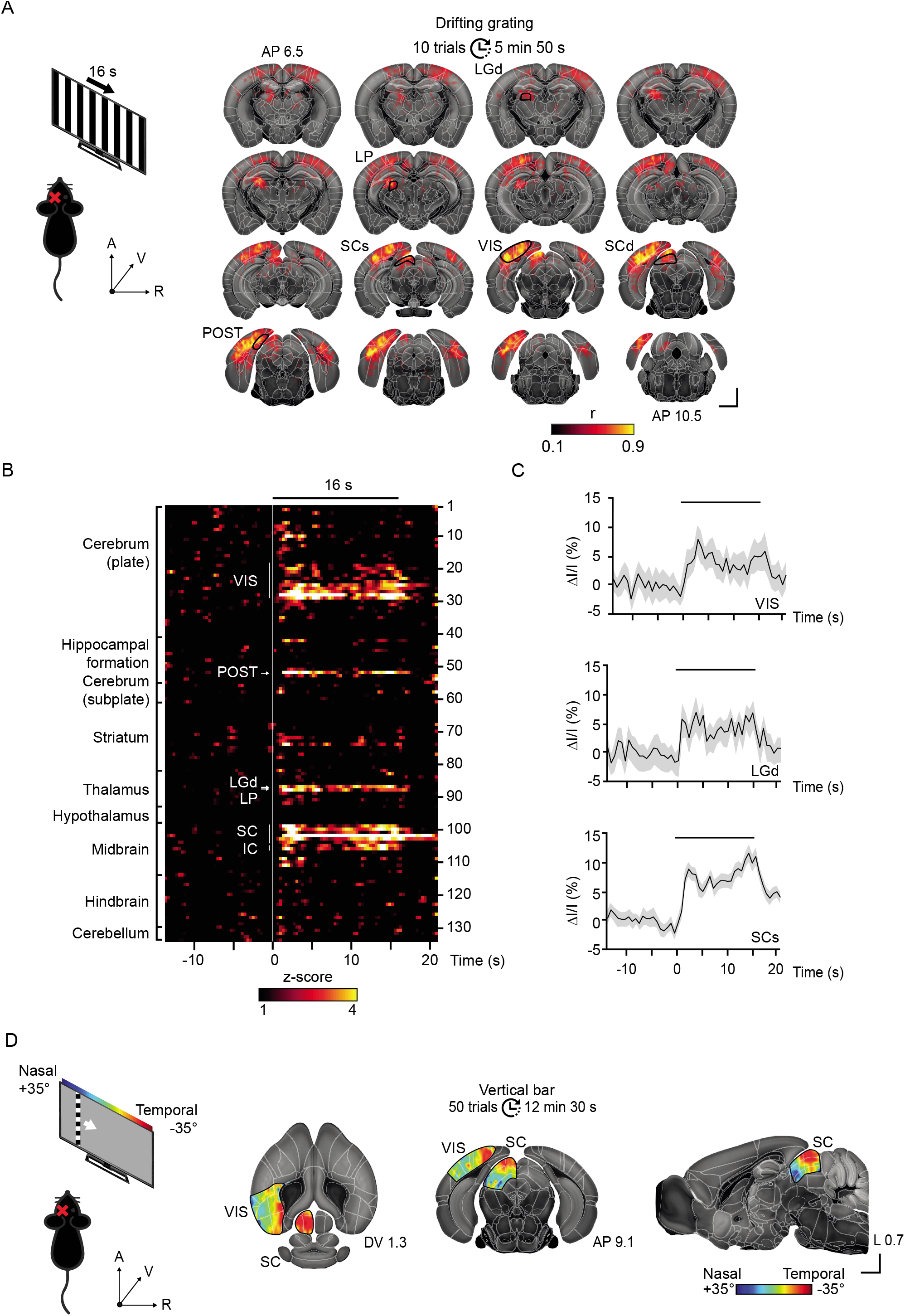
Brain-wide vfUSI of visual-evoked activity. **(A)** Left panel: schematic representation of the drifting gratings experiment for activity mapping of the visual system in response to the presentation of full-field vertical grating displayed during 16 s in the naso-temporal direction. The left eye was occluded (red cross). Right panel: example slices from the 3D correlation map presented in coronal view showing activated voxels during visual stimulation (average of 10 trials). Brain regions showing a statistically significant correlation to the stimulus are indicated in black (LP, LGd, SC, IC, VIS et POST) and outlined in the coronal sections. Scale bars, 1 mm. Significant activation corresponds to r values > 0.2. **(B)** Average hemodynamic responses (z-score) of the 134 regions in the right hemisphere. Regions were ordered by major anatomical structures (**Table S1**). Brain regions showing a statistically significant increase of hemodynamic activity are presented in white. Black thick line, stimulus. **(C)** Average hemodynamic response curves of VIS, LGd, and SCs brain regions presented in **(B)** Black thick line, stimulus. Grey highlighted traces, standard error of the mean. **(D)** Left panel: schematic representation of the experiments for retinotopic mapping along the azimuth axis. A horizontal bar (20° wide) was swept across a display monitor in front of the right eye (4°/s). The left eye was occluded (red cross). Right panel: example slices from a 3D retinotopic map in 3 different views crossing the visual cortex and superior colliculus superimposed on the anatomical atlas (greyscale). SC and VIS are outlined in black. Scale bars, 1 mm. A, anterior; P, posterior; D, dorsal; V, ventral; L, left; R, right. The horizontal bar indicates the timing of the stimulus. All color-coded maps are superimposed on Allen Mouse Brain atlas histological slices and white-outlined segmented brain regions.

### Functional mapping combining vfUSI and optogenetic stimulation

Similarly to previous opto-fMRI study (Desai et al., 2011), we combined optogenetic stimulation with vfUSI (termed opto-vfUSI) to identify both targets and circuit dynamics during activation of specific cell types. For the proof of concept, channelrhodopsin-2 (ChR2) activation of defined neuronal populations located in the SCs was performed, since it is a well-studied region playing an essential role in visuo-sensory processing (Oliveira and Yonehara, 2018) and involving long-range cortical and subcortical networks (**Figure 4A**). We used a Cre-dependent ChR2-eYFP reporter line crossed with several transgenic lines expressing Cre in either excitatory narrow-field cells (Grp-Cre) and wide-field cells (Ntsr-Cre) or inhibitory horizontal cells (Gad2-Cre; Gale and Murphy, 2014). Control experiments were performed by stimulation of the dorsal part of the retrosplenial cortex (RSP) located just above the SCs using the Thy1-ChR2 transgenic line expressing ChR2 broadly in pyramidal cells (**Figure 4A**). A short and low-intensity stimulation (40 repetitions, 2 ms pulse width, 20 Hz, 0.3 mW mean power) of either specific cell-types in transgenic mice or RSP in control mice elicited robust and reliable hemodynamic responses in the ipsilateral hemisphere in the activated region but also many areas distributed across the brain (**Figure 4B**). Activities in the contralateral hemisphere were not accessible due to implanted optical fiber (**Figure S1C; STAR Methods**).

**Figure 4.**
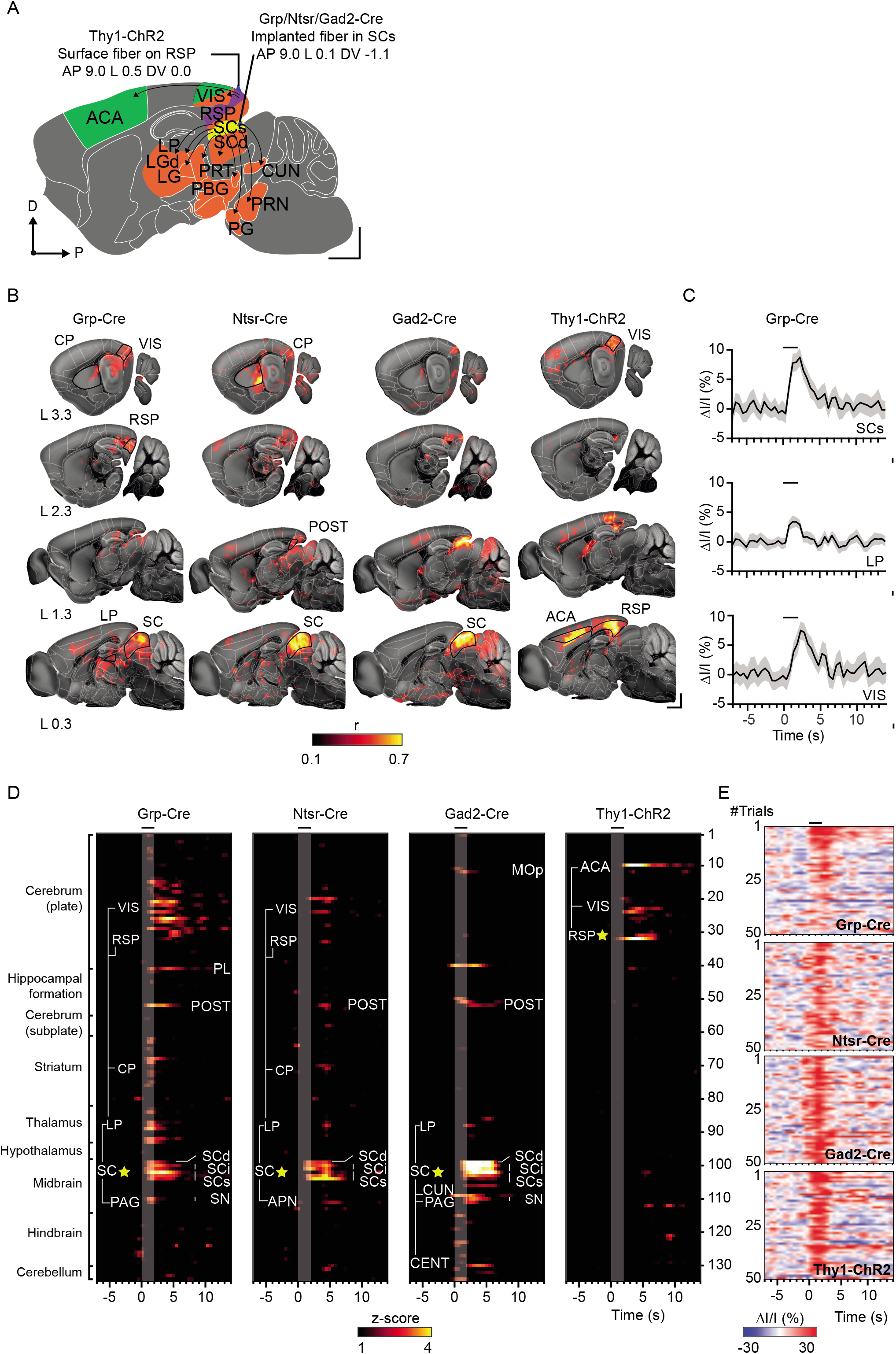
Functional mapping combining vfUSI and optogenetic stimulation. **(A)** Graphical representation of the main regions connected to the superficial layer of the superior colliculus (SCs). Arrows represent mono-synaptic connections between the SCs (highlighted in yellow) and other regions (highlighted in orange). The selected set of regions connected to the RSP (in purple) are highlighted in green. For optogenetic experiments, a fiber-optic cannula was implanted in the SCs for Grp, Ntrs, and Gad2 mouse lines. For control experiments, the fiber optic was positioned on the surface of the RSP in the Thy1-ChR2 mouse line. Scale bars, 1 mm. **(B)** Example slices from the 3D correlation map presented in the sagittal view showing activated voxels during optogenetic stimulation in Grp, Ntsr, Gad2 and Thy1-ChR2 lines (average of 50 trials for Ntsr, Gad and Thy1-ChR2 and 15 trials for Grp). Brain regions significantly activated are outlined in black in the sagittal sections. Scale bars, 1 mm. Significant activation corresponds to r values > 0.2. **(C)** Average hemodynamic response curve of SCs, LP, and VIS brain regions presented in **(D)** Black thick line, stimulus. Grey highlighted traces, standard error of the mean. **(D)** Average hemodynamic responses (z-score) of the 134 regions in the left hemisphere in response to optogenetic stimulation. Regions were ordered by major anatomical structures (**Table S1**). A selected set of brain regions showing a statistically significant increase of hemodynamic activity are presented in white. Black thick line, stimulus. Yellow stars indicate the location of the fiber-optic. **(E)** Trial-to-trial evolution of the hemodynamic response during optogenetic stimulation in each mouse line. Black thick line, stimulus. A, anterior; P, posterior; D, dorsal; V, ventral; L, left; R, right All color-coded correlation maps are superimposed on Allen Mouse Brain Atlas histological slices and white-outlined segmented brain regions.

Importantly, the circuits activated in Grp/Ntsr/Gad2-Cre lines were mostly overlapping. This included brain regions that have been reported to be anatomically connected to SCs through mono-synaptic projections, such as the intermediate (SCi) and deep (SCd) layers of the SC (Mooney et al., 1988) but also the lateral posterior (LP) nucleus of the thalamus (**Figure 4D**; Gale and Murphy, 2014). We also found differences between transgenic lines regarding regions that have been described as connected to SCs through di-synaptic circuits. Hence, the VIS was more broadly activated in Grp compared to Ntsr and even totally absent in Gad2 mice. Also, the cuneiform nucleus (CUN) and the IC were only activated in Gad2 mice, two targets of ipsilateral descending projections relaying behavioral function (García Del Caño et al., 2006; Redgrave et al., 1987) that are not activated in either Grp or Ntsr-Cre lines. Additionally, the Gad2 mice exhibited a high level of activity in the central lobule (CENT) of the cerebellum, known as part of the circuit involving the periaqueductal grey (PAG) which is playing a critical role in autonomous functions, motivated and threatening behavior (Deng et al., 2016; Faull et al., 2019). In the Ntsr mice, we observed activation in the anterior pretectal nucleus (APN) involved in various visual and oculomotor tasks (Prochnow and Schmidt, 2004) and the POST considered as an input structure to the hippocampal formation that relays visual landmarks coming from the RSP (Boccara et al., 2010; Kononenko and Witter, 2012). Consistent with the anatomical connectivity, RSP activation triggered an increase of the hemodynamic signal in all connected brain regions such as the VIS and the anterior cingulate cortex area (ACA; Mitchell et al., 2018), which strongly differs from patterns of activity observed in the other transgenic lines.

We observed that CBV responses to optogenetic stimulation measured both in cortical and subcortical regions fit well the canonical hemodynamic response function observed during evoked responses to short stimuli as previously observed in rats (**Figure 4C**; Urban et al., 2014b). Moreover, we noticed that onset time, peak amplitude, and post-stimulus decay obtained with opto-vfUSI (data not shown) are in the range of those measured in opto-fMRI (Desai et al., 2011). Additionally, opto-vfUSI revealed that repeated optogenetic stimulation of the SCs in the Grp mice led to a prolonged reduction in evoked hemodynamic response along the activated circuit (50% in 15 trials, **Figure 4E**), which was not observed in any other lines.

### vfUSI during an active sensory-motor task

Our last aim was to identify active circuits at a large scale during a task involving arousal, sensory, and motor components. We used water-deprived mice trained to reach for a water droplet with one paw after the presentation of a visual cue (3 s, 5-Hz blinking LED, left side; **Figure 5A**). Video recordings (**Video S2**) showed that trained animals performed the task with stereotyped movement kinematics within ~4 s after droplet delivery with a success rate comprised between 41 and 55% (**Figure 5C; STAR Methods**). We also observed that mice are moving both upper and lower limbs for several seconds before returning to a quieter state.

**Figure 5.**
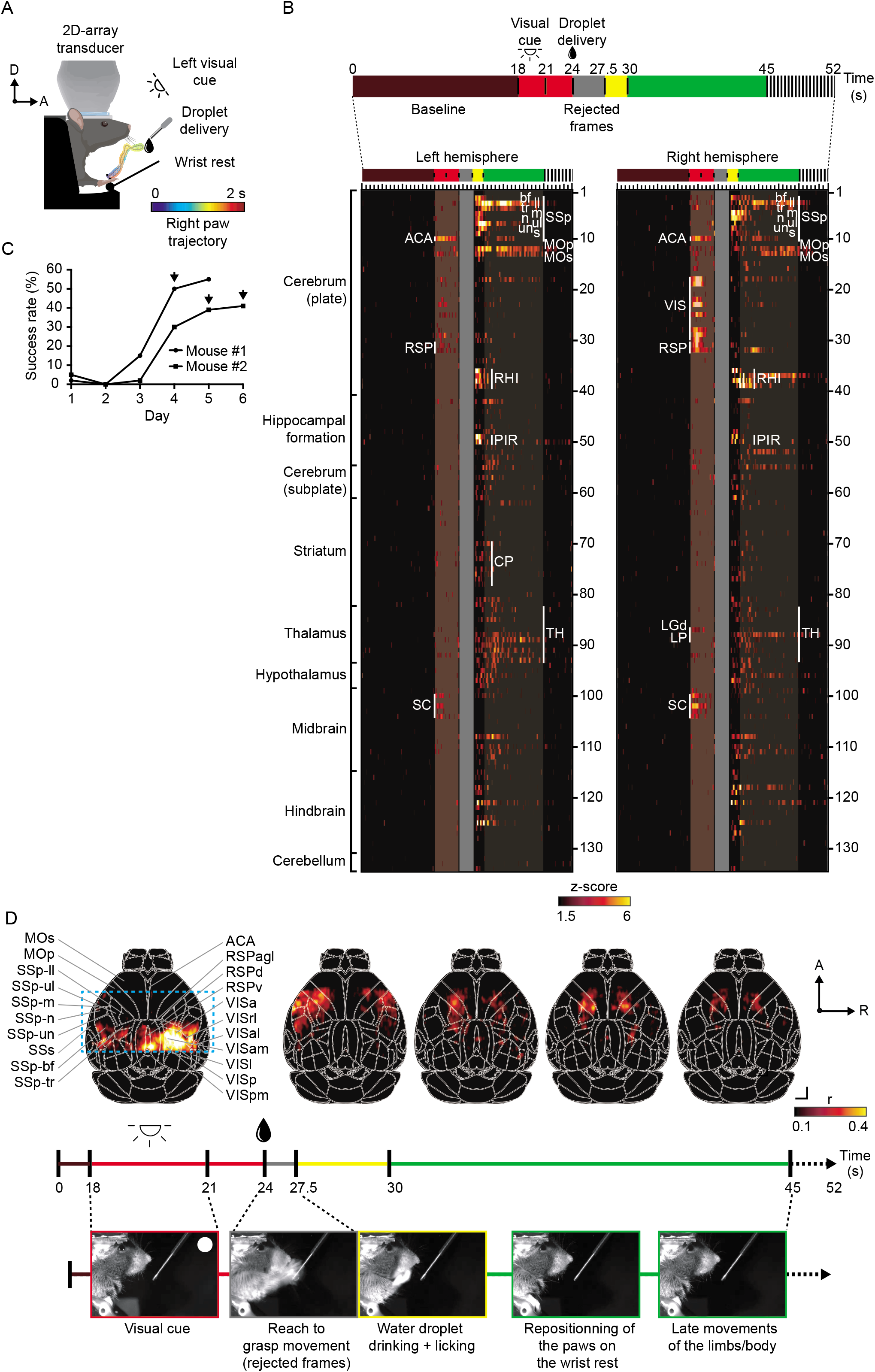
vfUSI during a sensory-motor task. **(A)** Schematic representation of the experimental design for vfUSI in head-fixed mice performing a reach to grasp droplet task. The position of the paw during the task for one typical trial is presented in rainbow color. A, anterior; D, dorsal. **(B)** Average hemodynamic responses (z-score) of the 134 regions in each hemisphere during the water droplet reaching task. Regions were ordered by major anatomical structures (**Table S1**). A selected set of brain regions showing a statistically significant increase of hemodynamic activity is presented in white. The top bar summarized the different phases during the task including i) the baseline, ii) the visual stimulation (red bar), iii) the reach to grasp task triggering strong artifacts in the vfUSI signal that was discarded from the analysis (grey bar, rejected frames), vi) quickly after the water droplet grasping (yellow bar), v) during repositioning of the paws on the wrist rest and additional limbs movements (green bar) and vi) the return to the quiet state (black dashed line). **(C)** Evolution of the water droplet reaching task success rate (%) to perform over experimental sessions. Black arrows indicate vfUSI data used for temporal analysis of the task. **(D)** Top view representation of hemodynamic activity in the cortex during the sensory-motor task. Top view color-coded correlation maps (top panel; r) in response to a visual cue (18-21 s) and motor task performance (27.5-45 s; left to right) with corresponding video frames of the animal performing the water droplet reaching task (bottom panel, left to right). Color-coded correlation maps are superimposed on white-outlined segmented brain regions. A list of abbreviations is provided in **Table S1**. A, anterior; R, right. Scale bar, 1 mm.

For the first time, we showed that temporal analysis of the hemodynamics across the brain has a complex pattern of activity involving both cortical and subcortical regions in both hemispheres (**Figure 5B**, n=45 trials). During the presentation of the visual stimulus, significant activation of the visual pathway was observed in the contralateral hemisphere (right), including VIS, LGd, LP, and SCs/i/d regions (**Figure 5B**, red bar, 18-24 s). Additionally, a marked bilateral activity was noted in several brain regions including the ACA, which is involved in higher-level functions such as attention allocation, reward anticipation and/or decision-making (Bush et al., 2002; Pardo et al., 1990), and the RSP, also known to play a role in diverse cognitive functions, including spatial navigation, orienting, memory and planning (Mitchell et al., 2018). During the initiation of the task and up to 3.5 s after droplet delivery, we observed strong artifacts due to brain movements, which are related to the fast movements of the limbs and the body. Consequently, all the corresponding frames were rejected (**Figure 5B**, grey bar, 24-27.5 s). Nevertheless, in such difficult imaging conditions, we took advantage of slow hemodynamic response decay time (~2 s) and the excellent temporal resolution (6 Hz) of vfUSI to analyze activity changes right after completion of the reaching task. During a short period (~2.5 s), we detected an increase of activity in the primary motor cortex (MOp) contralaterally to the right paw used for the task. Additionally, we found increases in the primary somatosensory cortex, particularly pronounced in the left upper limb (SSp-ul), the lower limb (SSp-ll), and, shortly, in the nose (SSp-n), mouth (SSp-m) and SSp-bf cortices used when the animal is drinking the droplet. Interestingly, a strong activity was detected in the perirhinal and ectorhinal cortices (RHI), which are involved in polysynaptic activation of the hippocampus by olfactory inputs (Liu and Bilkey, 1997) and the piriform (PIR) cortex. These results are consistent with recent experiments showing that head-fixed mice use chemosensory cues and olfaction to detect and spatially localize the presence of water (**Figure 5B**, yellow bar, 27.5-30 s; Galiñanes et al., 2018). In both hemispheres, we showed a prolonged activity correlated with repositioning of the animal for up to 20 s after water delivery in both SSp-ll and -tr (trunk) and primary and secondary motor cortices (MOp/s; **Figure 5B**, green bar, 30-45 s). Moreover, we observed activation in multiple thalamic regions (TH) which are relays of the motor and sensory information and in several parts of the caudoputamen (CP) including the ventrolateral area, well known to participate in the control of orofacial movements and forepaw usage accompanying feeding behavior (Pisa, 1988) and also in several nuclei of the reticular activating system (RAS) including the gigantocellular reticular nucleus (GRN) playing a crucial role in maintaining behavioral arousal (Gao et al., 2019).

A more detailed analysis of the spatial distribution of cortical responses (**Figure 5D; Video S2**) confirmed the sequential activation of visual and motor cortices contralaterally to the visual stimulus and the paw, respectively. Quickly after the movement of the paw, we observed the motor activity centered on MOp, in several SSp regions (mm, ul, ll, up, n, bf) and also in MOs. Several seconds after the task, the vfUSI signal was reduced mainly in the SSp cortices but was stably centered on the MOp. The activity observed in MOp, in both hemispheres, after the task was linked to limb movements during the repositioning of the animal on the wrist rest and are not associated with the droplet grasping.

## DISCUSSION

To the best of our knowledge, this is the first demonstration of volumetric functional ultrasound imaging of brain activity at a large scale in behaving head-fixed mice. Our goal was to develop a versatile hardware and software pipeline with advanced capabilities as compared to the initial proof of concept study performed in anesthetized rats (Rabut et al., 2019). Hence, we proved the high sensitivity of the vfUSI platform and showed that whisker-evoked hemodynamic responses are detectable in the cortex and the thalamic relay after an experimental session of 30 minutes maximum. Reducing the overall duration of an experiment and the number of stimuli is crucial to preserve proper animal physiology, animal comfort, and avoid the effects of habituation (Miller et al., 2018). Even if vfUSI has a voxel size ~2.5 times lower (~250 μm) compared to 2D fUSI (**Figure S4**), we were able to detect hemodynamic signals resulting from neuronal activation of functional units such as small individual cortical columns (~100 μm) during stimulation of a single whisker, or microcircuits processing orientation and directional selectivity during visual stimulation.

Regarding the temporal resolution of vfUSI, the framerate could be adapted on-demand from 1.4 to up to 6 Hz by decreasing the number of compound images with little impacts on sensitivity. The range of use is appropriate for imaging hemodynamic responses that are relatively slow because these are driven by a complex nonlinear function depending on various neuronal and vascular changes (Iadecola, 2017). Importantly, vfUSI can be combined with acute or chronic electrophysiological recordings to address the limitation of slow blood flow dynamics offering multimodal capabilities across various spatiotemporal scales, as was previously demonstrated with 2D fUSI (Macé et al., 2018; Urban et al., 2014c).

We showed that opto-vfUSI is a new method to identify cortical and subcortical targets recruited by optogenetic activation of specific cell populations, offering repeated assessment, potentially over extended periods suitable for longitudinal studies of brain function. Note that a previous study combining fUSI and optogenetics reported an energy-dependent photodilation in naïve mice using various light intensities (from 2 to 40 mW.mm^-2^; Rungta et al., 2017). It suggests that the light stimulation could have direct effects on the vasculature, which should be taken into consideration when combining hemodynamics imaging methods such as fMRI or fUSI with optogenetics. To minimize such potential effects, in our experiments, we used a lower light intensity (1.5 mW.mm^-2^) and did not observe any significant increase of the hemodynamic signal in control experiments in WT mice in these conditions. In one transgenic line, we observed a rapid decrease in the evoked hemodynamic response after only a few stimulus repetitions that only recovered several hours later, which may indicate a potential habituation or deactivation of brain circuits. Another hypothesis is that weak, modulatory functional or transient connections may not be detectable because of the slow amplitude of the hemodynamic signal as compared to baseline fluctuations. Overall, these results suggest that the effects of localized optogenetic manipulation of neurons could lead to misinterpretation as recently reported (Li et al., 2019). Nevertheless, opto-vfUSI remains a tool of choice for *in vivo* circuit mapping during modulation of genetically-defined neurons.

For the first time, we identified the specific dynamics of brain activity during a water droplet grasping task sequentially involving arousal, visual, motor, and sensory circuits. However, we were not able to image activity during the entire task due to artifacts produced by strong limb movements. Implantation of a lightweight and miniature 2D-array transducer directly on the head similar to what we developed for fUSI in freely-moving rats (Urban et al., 2015) is a potential solution. For this purpose, micro-machined ultrasonic transducers (MUTs) may be the most relevant technology (Oralkan et al., 2003). This strategy may overcome the limitations of the head-fixed configuration showing a reduced neuronal activity in the sensory cortex (De Kock and Sakmann, 2009) and allow vfUSI in freely-moving rodents.

For the sake of conciseness, detailed results on vfUSI for mapping spontaneous resting-state functional connectivity (RSFC) are not presented here. Nevertheless, this application could be of high relevance to study the functional connections of specific brain regions and local circuits, as well as important new insights in the overall organization of functional communication in the brain circuit (van den Heuvel and Hulshoff Pol, 2010). Here, we devised a similar approach to the one described in the 4D fUSI paper in anesthetized rats (Rabut et al., 2019) using seed-based (**Figure S3A**) and correlation matrix analysis (**Figure S3B**, and **Table S2**). We observed a high degree of connectivity between mono-synaptically connected regions with robust isocortical, striatal, thalamic, and hippocampal circuits that could be observed in awake mice. Additionally, we showed that awake mice display reinforced connectivity involving thalamo-cortical pathways as compared to available literature, which was possibly influenced by the effect of the anesthesia (Grandjean et al., 2017). vfUSI is a credible alternative to BOLD-fMRI for functional connectivity studies in preclinical research while offering a better spatiotemporal resolution, versatility, ease of use, and affordability.

## Supporting information

Supplementary code 1 - Registration

Supplementary code 2 - Retinotopic mapping

Supplementary code 3 - Resting state functional connectivity

## ACKNOWLEDGMENTS

This work was supported by grants from the Leducq Foundation (15CVD0_2_), from FWO (MEDI-RESCU2-AKUL/17/049, G091719N, and 1197818N), from VIB TechWatch (fUSI-MICE) and from internal NERF TechDev fund (3D-fUSI project). We thank Dr. M. Krumin for assistance with the design of the mouse holder and Dr. C. Cowan for help with the Blender software for 3D rendering. We also thank the NERF Pls for proofreading of the manuscript and all NERF animal caretakers, including I. Eyckmans, F. Ooms, and S. Luijten, for their help with the management of the mouse lines.

## AUTHOR CONTRIBUTIONS

Development of the hardware and the software was performed by A.U. and G.M.

Animal preparation and experiments were performed by C.B., M.G., G.M., T.L., A.S-D. and A.U. Data analytics were carried out by A.U., C.B., M.G., G.M., T.L. A.S-D. and E.M.

The figures and schematics were designed by A.U. and prepared by C.B., M.G., T.L. and G.M. Finally, A.U. and G.M. wrote the manuscript with recommendations and corrections from all authors.

## STAR*METHODS

### RESOURCE AVAILABILITY

#### Lead Contact

Further information and requests for resources and reagents should be directed to and will be fulfilled by the Lead Contact, Alan Urban.

#### Materials Availability

This study did not generate new unique reagents.

#### Data and Code Availability

The published article includes all code generated or analyzed during this study (see **Supplementary_codes-vfUSI**).

## EXPERIMENTAL MODEL AND SUBJECT DETAILS

### Mice models

Adult male C57BL/6J WT mice (KU Leuven, Belgium) were used for whisker (n=3), visual stimulation (n=2), resting-state functional connectivity (n=3), and behavioral task experiments (n=3). Progeny from Tg(Ntsr1-cre)GN209Gsat/Mmucd (Ntsr) (MMRRC, USA), Gad2^tm2(cre)zjh^/J (Gad2) (The Jackson Laboratory, USA) and B6.FVB(Cg)-Tg(Grp-cre)KH288Gsat/Mmucd (Grp) (MMRRC, USA) lines respectively crossed with Gt(ROSA)26Sor^tm32(CAG-COP4*H134R/EYFP)Hze^ (ChR2) (The Jackson Laboratory, USA) and B6.Cg-Tg(Thy1-COP4/EYFP)18Gfng/J (Thy1) (The Jackson Laboratory, USA) lines (n=2, 3, 3, and 1, respectively) were used for the optogenetic experiments. All mice weighed between 20–30 g.

### Rat model

Adult male Sprague-Dawley rats weighing between 250–350 g (n=4; Janvier Labs, France) were used for whisker stimulation experiments.

Rats and mice were kept in separate rooms following a 12-hr dark-light cycle with a constant temperature of 23 °C and *ad libitum* access to food and water. After surgery, each rodent was housed alone in its cage. Imaging sessions were performed in separate rooms dedicated to either mouse or rat. To reduce the stress, rodents were handled and habituated to the experiment room environment 1 to 2 weeks before the experiment. The experimental procedures were approved by the Committee on Animal Care of the Katholieke Universiteit Leuven, following the national guidelines on the use of laboratory animals and the European Union Directive for animal experiments (2010/63/EU).

## METHOD DETAILS

### Cranial window procedures

For induction, mice were anesthetized with isoflurane (3% for induction; Iso-Vet, 1000 mg/g, Dechra) in room air and maintained with ketamine/medetomidine (Nimatek 40 mg/kg, EuroVet and Domitor 1 mg/kg, Vetoquinol; respectively). Once the scalp was removed and skull cleared from tissue, a stainless steel custom designed head positioner was fixed to the animal skull with dental cement. A cranial window was performed with care to avoid impairment of the dura mater. In WT mice, the cranial window extended from bregma +3.0 to −4.5 mm and ± 5.0 mm bilaterally. The window was designed for vfUSI with 2-3 sectors of the 2D-transducer array in coronal direction (**Figure S2**).

For optogenetic experiments, a cranial window was opened from bregma +3.0 to −6.5 mm and laterally from +5.5 to −1 mm over the left hemisphere. In Ntsr, Gad2, and Grp transgenic lines, a 200-μm diameter fiber-optic cannula (OGKL2-1, Thorlabs) was implanted in the superior colliculus (coordinates tip of the fiber: AP=9.0 L=0 V=-1.1 mm) and tilted at 54°. The cannula was then fixed with dental cement. The window enabled to image with two sectors of the probe in the sagittal direction (**Figure S2**).

Rats were anesthetized with isoflurane (5% for induction, 2% for maintenance; Iso-Vet, 1000 mg/g, Dechra) and then fixed in a stereotaxic frame. After scalp removal and tissue cleaning, a stainless steel custom designed head-positioner was fixed with screws and dental cement to the animal skull. An ~11-mm^2^ cranial window was opened from bregma +4.0 to −7.0 mm and 6.0 mm apart from the sagittal suture. The dura was kept intact. This window was imaged using 3 or 4 sectors of the probe in coronal direction (**Figure S2**).

In all animals, a silicone elastomer layer (Body Double-Fast set, Smooth-On, Inc, USA) was used to seal the cranial window. An additional head shield was added on top of the head-positioner as a protection (Urban et al., 2015). Both mice and rats were placed in a warm cage directly after surgery and monitored until they woke up. Analgesic (Buprenorphine, 0.1 mg/kg, Ceva), antibiotic (Cefazoline, 300 mg/kg, Sandoz) and anti-inflammatory (Dexamethasone, 0.5 mg/kg, Dechra) drugs were injected 24 and 48 h after the surgery. A second antibiotic (Emdotrim, 1 %, Ecuphar) was added to the water bottle.

### Head fixation and imaging session

After five days recovery, mice were trained to be head-fixed while awake by screwing the head-positioner to the animal holder. Rats were shortly anesthetized with 2% isoflurane during installation in a suit (Martin et al., 2006) and fixation of the head before waking up. For both models, the period of fixation was progressively increased from 5 min to 2 h. A 5% sucrose solution (Sigma-Aldrich, USA) was given to the animals along with the training session as a reward.

Before the imaging session, the silicone cap was removed and replaced by a thin layer of 2.5% agarose solution (Sigma-Aldrich, USA) covered with ultrasonic gel. Then, the ultrasound probe was placed and adjusted over the imaging window. After the imaging session, the agarose layer was removed and replaced by a silicone layer.

### Whisker stimulation

For both mice and rats, whiskers’ stimulations were delivered using a piezoelectric actuator (BA6020, PiezoDrive, Australia). A comb fixed to the actuator was used for multiple whisker stimuli whereas, a capillary glass of 500 μm diameter (Sutter, USA) was used for individual whiskers. The stimulus was a 5-Hz sinusoidal deflection of ~20° of amplitude delivered for 6 s (5 s for rats). Each trial consisted of 14 s baseline, 6 s stimulus, and 15 s recovery for a total of 50 images.

### Visual stimulation

Visual stimuli were presented in a 28’’ monitor (llyama, USA) placed in landscape orientation 18 cm away from the right eye and 45° from the anteroposterior axis of the mouse. The background color was set to 50% grey luminosity. The sight from the left eye was blocked with a small piece of paper. The grid stimulus was a full field vertical drifting grating (black and white) of 20° spatial frequency moving in nasal to the temporal direction at 10°/s. Each trial consisted of 20 s baseline, 16 s stimulus, and 5 s recovery for a total of 50 images.

The retinotopic stimulus consisted of a 15° wide vertical bar moving in naso-temporal direction at 4°/s. The bar was filled with a black and white checkboard pattern (25° spatial frequency) reversing at 6 Hz. Each trial consisted of a 2.1 s baseline and 14 s stimulus for a total of 20 images.

### Optogenetic stimulation

Optogenetic stimulation consisted of 40 pulses of 2 ms at 20 Hz using a 473-nm DPSS laser (R471003GX, Laserglow Technologies, USA). The average power at the tip of the fiber-optic was 0.3 mW (~1.5 mW/mm^2^). For the non-implanted Thy1 mice, a cannula was positioned with a micromanipulator on the surface of the retrosplenial cortex at coordinates AP=9.0 L=0.5 V=0 mm. Each trial consisted of 14 s baseline, 2 s stimulus, and 19 s recovery for a total of 50 images.

### Sensory-motor task

The water droplet reaching task protocol was adapted from Galiñanes et al., 2018. The training phase occurred in a transparent chamber of 10×10×10 cm^3^ with a vertical opening of 9×20 mm^2^ placed in the center of one wall giving access to a tube for water delivery placed outside the chamber. A 3-s visual cue (5-Hz blinking) was triggered 6 s before delivering a single 5-μl water droplet. After recovery, mice were placed under water restriction (1 ml/day) and trained, while freely moving inside the chamber to reach and catch the water droplet with the forepaw. Once a 70% success rate was reached, mice were trained to perform the same task in head-fixed conditions in the vfUSI imaging set up. Each trial lasted 50 s and was imaged at 6 Hz (300 frames). Visual cue started at frame 100 (16.6 s), and the droplet was delivered at frame 136 (22.6 s). Animal movements were recorded using an infrared camera at 100 frames per second (GC660, Allied Vision, USA) and analyzed manually.

### Resting-state functional connectivity

After head-fixation, the animal was let to recover from manipulation for 30 min before starting the imaging session. The animal was imaged in resting conditions in the dark continuously during one h at a vfUSI imaging framerate of 0.8 images per second (4500 images). For the analysis of restingstate functional connectivity data, we used a seed-based and correlation matrix approach, as described in Rabut et al., 2019.

### Cytochrome oxidase staining

After the experiment, mice received transcardial perfusion with 4% PFA, and the brain was collected and fixed for 24 h in 4% PFA before being placed in PBS. Flattened cortices were prepared following the cytochrome oxidase staining protocol described by Lauer *et al*., 2018 (Lauer et al., 2018). Briefly, after separating the two hemispheres and removing the subcortical structures with a razor blade, the hemispheres were flattened between two glass slides for 4-5 h at 4°C. The flattened hemispheres were then post-fixed in 4% PFA and stained with the cytochrome oxidase staining solution made with HEPES, sucrose, nickel-ammonium sulfate (NiAS), cytochrome C and diaminobenzidine (DAB) for several hours. The stained slides were imaged with a microscope at 20x magnification (Olympus BX51WI, USA) equipped with an IR camera (RETIGA 2000R, Qimaging, USA). Data were post-processed using the Fiji package of ImageJ (NIH, USA).

### Ultrasound sequences

Different ultrasound sequences were used to perform vfUSI experiments. Each sequence is characterized by five parameters as follow:

*Sectors* Number of sectors used to image. Each sector covers a surface of 2.7×9.6 mm^2^.
*Averaging* Number of averaging used for a plane-wave acquisition.
*Angles* Set of angles used for producing a compound image.
*Ncompound* Number of compound images for producing a single Doppler image.
*Frate* Final framerate of the vfUSI acquisition.

A plane-wave image is acquired by sequential emission in all the *Sectors* (**Figure S1B**) and averaged several times (*Averaging*). A set of tilted plane waves with n *Angles* are emitted to create a compound image. Finally, a single Doppler image is computed using a set of *Ncompound* images. **Tables 1** and **2** summarize the different sequences used in each experiment.

### Computing of the 3D Doppler images

The data was digitalized in 14 bits at a 60 MHz sampling rate, but only half-band (8 to 25 MHz) was used by the transducer. To reduce the bandwidth by a factor 2 (from 4.4 to 2.2 GB/s), only two values were kept for all samples. The original data sampled at 60 MHz was recovered with an interpolation filter implemented directly at the GPU level.

A double buffer was implemented inside the processing electronics allowing simultaneous recording of the vfUSI data in one buffer while the second is transferred to the workstation. Such architecture allows continuous data transfer at an average rate of 2.2 GB/s. The processing of compound images composed of up to 1400 plane-wave images is distributed in 4 GPUs and performed as followed:

#### RF Data Interpolation

The subsampled data was interpolated to 60 MHz using a bandpass FIR filter of 41 coefficients between 25% and 75% of the band. A second low pass FIR filter of 25 coefficients at 25% of the band interpolated the sample to 240 MHz. This filter enabled a higher phase resolution during the beamforming procedure.

#### Beamforming

The raw ultrasound data is composed of the 1400 plane-wave emissions that were beamformed using a classical delay-and-sum algorithm adapted to 3D plane-wave imaging. The seven planewave images of each compound image were added to obtain 200 compounded images. The beamforming voxel was 220×280×175 μm^3^ which was approximately the resolution of the image.

#### Filtering

The beamforming output is a set of 200 3D compound images annotated:

*s*(*x,y,z,t*), *t* = 1...200.

These images are a superposition of signal coming from both the blood flow and the tissue:

*s*(*x,y,z,t*) = *s_Blood_*(*x,y,z,t*) + *s_Tissue_* (*x, y, z, t*).

As in the 2D fUSI, the blood component was separated from the tissue component by eliminating the first singular values of this data (Urban et al., 2005).

#### Doppler Image

The Doppler image is obtained as the intensity of the blood signal: 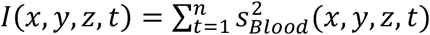. The 4 GPUs are parallelized with a delay of 1/4 of the computing period (**Figure S1B**). Using this parallelization of acquisition, data transfer, and computing, an average bandwidth of 2.2 GB/s was obtained corresponding to a vfUSI period of 0.7 s for acquiring each image (0.5 s acquisition, 0.2 s dead time, 71% duty cycle). By reducing the number of angles to 3, a 100% duty cycle is reached (see sequence S2fast in **Table 1**).

#### Activity

The activity is defined as the relative increase of the intensity image compared with a baseline

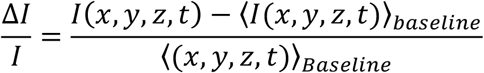

in where 〈 〉 indicates the temporal mean.

## QUANTIFICATION AND STATISTICAL ANALYSIS

### Registration and segmentation

In order to facilitate inter-animals comparisons and provide a general readout of the brain activity, we designed a registration/segmentation software adapted to vfUSI data (**Code S1**). It consists of 4 parts:

#### Atlas

The registration and segmentation are based on the Allen Mouse Common Coordinate Framework (CCF, v3; Oh et al., 2014). The digital atlas is segmented in 622 regions with a 25-μm^3^ voxel resolution. Data size and processing time were reduced by subsampling the atlas to a 50-μm^3^ resolution. It was then adapted to vfUSI by eliminating non-vascular regions (ventricles and white matter). Small areas (<0.5 mm^3^) below the technology resolution (220×280×175-μm^3^ voxel size) were pooled with neighboring anatomical regions. Olfactory bulbs and posterior cerebellum areas were excluded from the list because not imaged in this paper. The laminar organization of the cortex, hippocampus, and superior colliculus were removed, and layers were pooled. Regions larger than 15 mm^3^ (e.g., caudoputamen) were anatomically subdivided. The list contains a set of 134 selected regions/hemisphere (including 220 individual regions in total; **Table S1**).

#### Registration

For each imaging session, a single Doppler volume was used for registration to the Allen Mouse Brain atlas. Such a volume was interpolated to 50-μm voxel size and semi-automatically registered using affine transforms performed through a graphical user interface (**Figure 1B**). The transformation was saved and systematically applied to the data set associated with the full experimental session. Following this procedure, all output data we used was registered to our custom Allen Mouse Brain atlas.

#### Display

Registered datasets (Doppler, Correlation maps, etc.) can be displayed in all coronal, axial, and sagittal views superimposed on the segmented brain atlas.

#### Segmentation

Registered datasets were segmented in 134 brain regions/hemisphere. For each Doppler image and time point of the imaging session, the averaged intensity of all voxel in the 134 regions over time were extracted.

### Resolution

vfUSI resolution has been measured by imaging a 150-μm bead (phantom) placed inside a block of 1.5% agarose. The bead was imaged with a 50 μm^3^ beamformed voxel. The Doppler image resolution was measured at 220×280×175-μm^3^, respectively, in the *x, y, z* dimensions (**Figure S1A**). The resolution improvement in the *x* dimension is related to compound angles.

### Activity signal

The activity signal is defined as the relative increase of the Doppler intensity for the baseline signal.

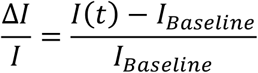

### Correlation map

The median value was calculated in all trials of a session. The median value was preferred as compared to the mean value to remove outliers linked to animal movements. The vfUSI images were computed as the Pearson’s correlation coefficient between the median images *I*(*x, y, z, t_t_*) and the temporal stimulus pattern *A*(*t_t_*), *i* = 1..*N*

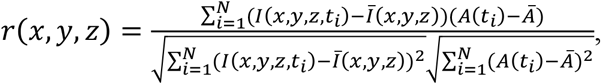

where 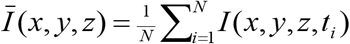 and 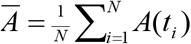. A median spatial filter of 3×3 pixels was applied to the final correlation map. The *z*-score value was then calculated using a Fisher’s transform, 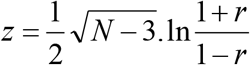. Only pixels with a z-score>1.6 were considered significantly activated (p<0.05 for a one-tailed test).

### Retinotopic mapping

The median value was calculated over all the trials in a session. To produce the retinotopic map, across-correlation between temporal signal *s*(*t*) and the maximal peak signal *h*(*t*) was computed for every voxel in the data. The peak signalwas set as a Hanning window of 7 points length (4.9 s). The time point associated with the maximum signal was localized by cross-correlation,directly matching the angular position of the displayed bar to the time point. Eventually, a spatial median filter of 3×3 pixels was applied to the final map (see **Code S2**).

### Processing of the resting-state functional connectivity data

The raw data set consisted of 4500 3D images sampled at 1.4 Hz. The following set of filters (see **Code S3**) was applied to extract the connectivity map:

#### Noise filter

Animal movements generated large artifacts in the vfUSI signal corresponding to high positive values. To filter such noise, the set of 4500 images was divided into 300 consecutive sets of 15 images from which we kept the data with the lowest mean intensity. This process resulted in a new set of 300 images that was further processed.

#### Frame rejection

Remaining noisy frames were eliminated by computing the average intensity of all 3D images and the associated standard deviation. Frames showing an intensity 1.8 times higher than the standard deviation were replaced by the previous one.

#### High-pass filter

A high-pass filter (Butterworth of order 2) with a cutoff frequency of 2-MHz was also applied.

#### Filter Average

Brain-wide variations in the vfUSI signal were filtered by subtracting the average intensity across the brain (*s*(*t*))to the signal in each voxel.

#### Normalization

The signal in each voxel was normalized by its mean intensity.

The filtered data were registered and segmented in the 134 brain regions of the Allen Mouse Brain Atlas to obtain a matrix *P*(*r, t*) where *r* is the region and *t* the time. Common brain-wide signals were discarded by removing the first singular value of the matrix *P*(*r, t*). Then, the correlation matrix *C_iJ_* was computed as a Pearson’s correlation coefficient between regions *r_i_* and *r_j_*.

### General processing of the segmented data

In visual and optogenetic experiments, several trials were repeated. The segmented data is represented as a 3D matrix *P*(*r, t, T*) where *r* is the region, *t* the time and *T* the trial. 3D matrix data were computed as the mean value in each region 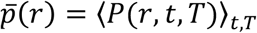 and outliers eliminated if 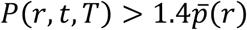. Then, the 3D matrix was converted into a z-score as follow: 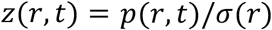, with 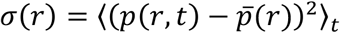.

Brain-wide variations in vfUSI signal were removed from *z*(*r, t*) using the Filter average as described above in the Resting-state functional connectivity section.

### Processing of the sensory-motor task

Individual trials were visualized and included in the analysis workflow as successful trials when the mouse performed the water droplet reaching task during the 4-s period following the drop delivery. The median value was calculated before applying the Filter Average as described in the restingstate functional connectivity section. Correlation maps were then computed for visual cue and different time windows of the task.

## VIDEOS TITLES

**Video S1. Top left panel**, 3D correlation map of multiple whisker stimulation (average of 5 trials); **Top right panel**, 3D correlation map of multiple whisker stimulation (average of 50 trials); **Bottom left panel**, 3D correlation map of individual whisker stimulation (average of 50 trials); **Bottom right panel**, 3D correlation map of visual stimulation (average of 10 trials). Active voxels are color-coded based on correlation (r; scale in **Figure 2B**, left)

**Video S2.** Top view representation of the cortical activity during the sensory-motor task (average of 45 trials). The left panel shows a typical video recording during execution of the task.

## SUPPLEMENTAL INFORMATION TITLES AND LEGENDS

**Figure S1** – 2D-array transducer spatial resolution, 3D ultrasound sequence and animal preparation used for vfUSI.

(A) Spatial Resolution of the 2D-array transducer. A 150-μm diameter bead was imaged inside a 1cm^3^ block of 1.5% agarose with a 50×50×50 μm^3^ beamforming voxel size. The Doppler image resolution was 220 μm in the azimuth (x), 175 μm in the elevation (y), and 280 μm in the axial (z) dimensions.

(B) vfUSI acquisition and real-time processing. Data was transferred to the workstation during the acquisition. All the data of a Doppler image were received and then dispatched to one GPU for processing. The 4 GPUs are working in parallel with a delay of 1/4 of the computing period. Data transfer and computing have a bandwidth of 2.2 GB/s for a vfUSI period of 0.7 s per frame (0.5 s acquisition, 0.2 s dead time, 71% duty cycle).

(C) Animal preparation. Top view schematic of the positioning of the cranial window and stainless-steel positioner for vfUSI in awake head-fixed mice (left and middle panels) and rats (right panel). In mice, the 2D-array transducer was placed in coronal (left) or in sagittal orientation (middle) to allow the fiber-optic cannula positioning for optogenetic experiments. In rats, bone screws were added to reinforce fixation of the head positioner, and the transducer was positioned in coronal orientation. A, anterior; V, ventral; R, right.

**Figure S2 - vfUSI during multiple or single whisker stimulation in awake head-fixed rats.**

Top panel. Correlation map during stimulation of multiple whiskers (25 deflections, 5 Hz, 5 s, average of 30 trials). Coronal sections are presented from anterior to posterior direction (top to bottom and left to right). The corresponding temporal hemodynamic responses and plots confirmed the stability across trials.

Bottom panel. Correlation map during stimulation of a single whisker (25 deflections, 5 Hz, 5 s, C3 in blue, Beta in green, and B1 in red). The number of trials is displayed at the bottom of each panel. Coronal sections are presented from anterior to posterior direction (top left to bottom right corner). Averaged hemodynamic response curves for C3, Beta, and B1 stimulation extracted from the correlation maps are presented below with the corresponding individual trial traces. Experiments were performed in the same rat. Vertical grey bar, the period of whisker stimulation. Scale bar, 1 mm. Grey highlighted traces, standard error of the mean. Color code correlation maps are superposed to the Doppler images. Significant activation corresponds to r values > 0.2.

**Figure S3 - Resting-state functional connectivity in awake mice.**

(A) Resting-state functional connectivity maps using seed-based correlation. Example slices of the correlation maps (r) superimposed on Allen Mouse Brain Atlas histological slices (top anterior to posterior bottom coronal planes) for cortical, hippocampal, hindbrain, and thalamic seeds. Scale bar, 1 mm.

(B) Connectivity matrix for the 134 brain regions/hemisphere. The horizontal axis corresponds to connectivity seeds of the right hemisphere. The left vertical axis corresponds to connectivity targets in the left hemisphere (bottom left of the matrix). The vertical right axis corresponds to connectivity targets in the right hemisphere (top right of the matrix). Regions are organized according to **Table S2**.

**Figure S4 - 2D-array vs. linear transducers**

Comparison of ultrasound probe specifications between the 2D-array and the linear transducer in the same mouse. 3D Doppler images were acquired with the 2D-transducer array (middle). A mechanical scan of the coronal cross-section was performed with the linear transducer array

(bottom). Scale bar, 1 mm.

**Table S1 -** List of the 134 brain regions/hemisphere classified by major anatomical structures adapted from the Allen Mouse Common Coordinate Framework.

**Table S2 -** List of the 134 brain regions/hemisphere classified by major anatomical structures adapted from the Allen Mouse Common Coordinate Framework used for resting-state functional connectivity experiments presented in **Figure S3.**

