## Supplemental File1 for "A platform for brain-wide functional ultrasound imaging and analysis of circuit dynamics in behaving mice"

Figure S1. 2D-array transducer spatial resolution, 3D ultrasound sequence and animal preparation used for vfUSI

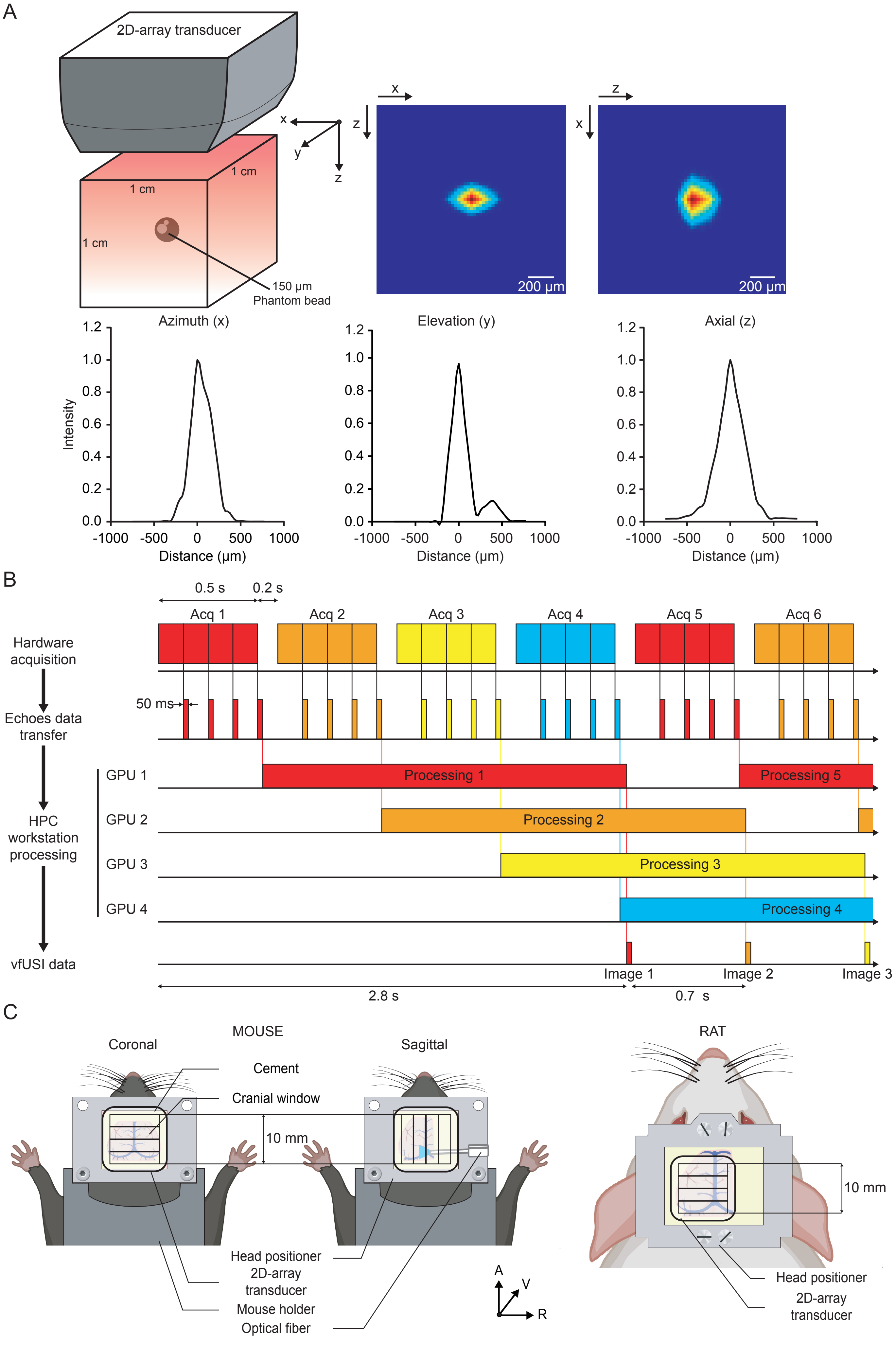

### Figure S2. vfUSI during multiple or single whisker stimulation in awake rats

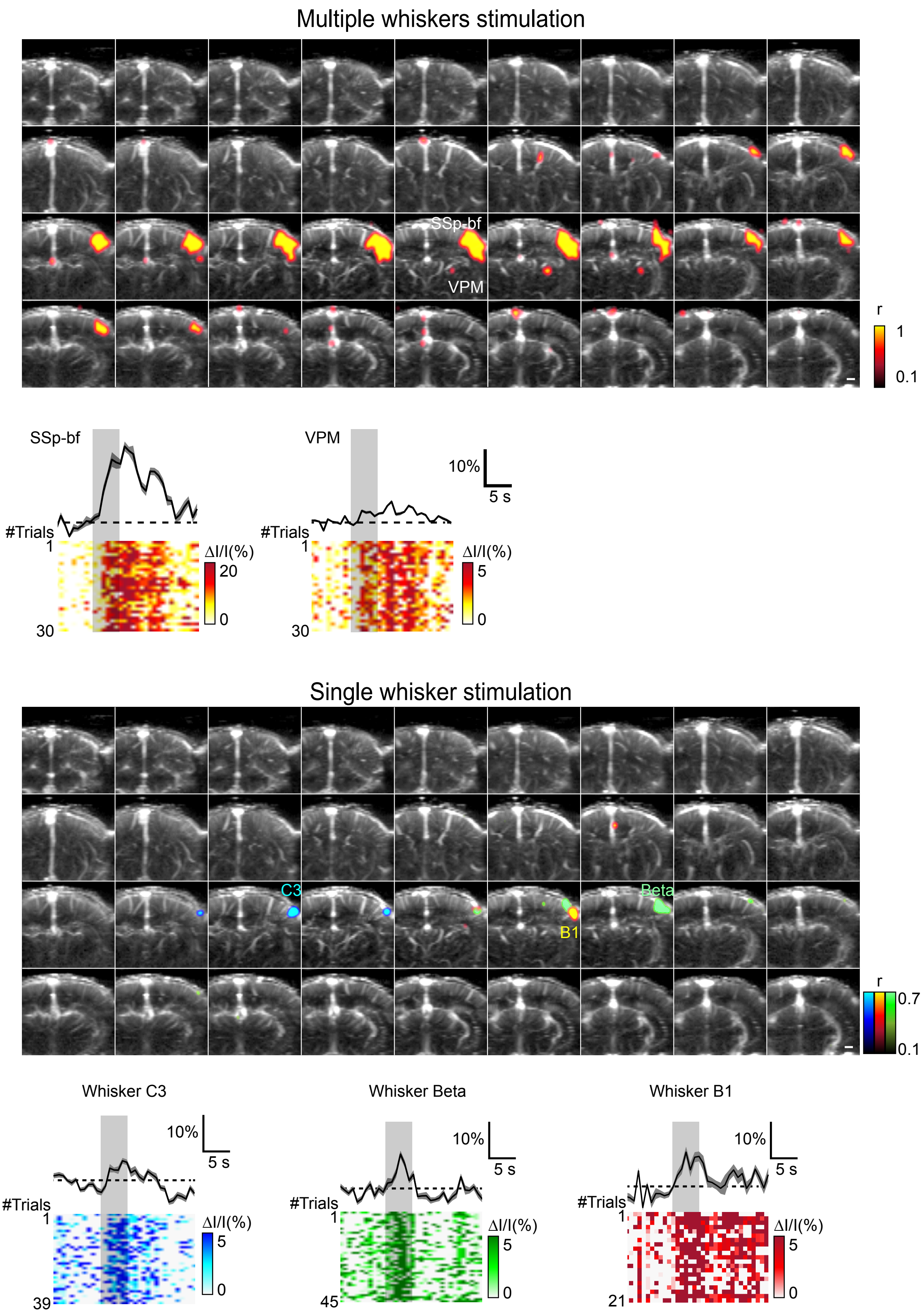

### Figure S3. vfUSI resting-state functional connectivity in awake mice

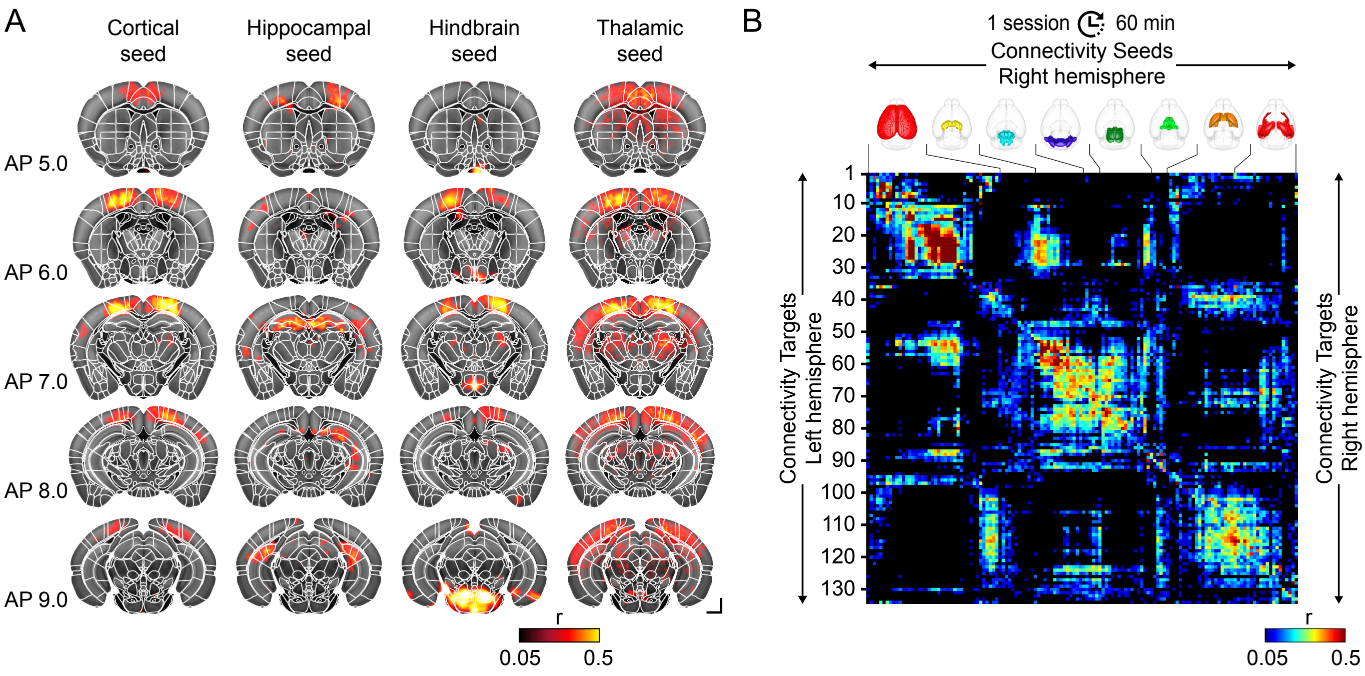

Figure S4. 2D-array transducer vs linear transducer

**2D-array transducer**
1024 piezo-elements - channels
Frequency: 15 MHz
Electronic focusing (no lens)
Voxel size: 220x280x175 $\mu m^3$
Field of view: 9.6x9.6x10 $mm^3$
Frame rate: 1.4-6 Hz
3D Doppler image

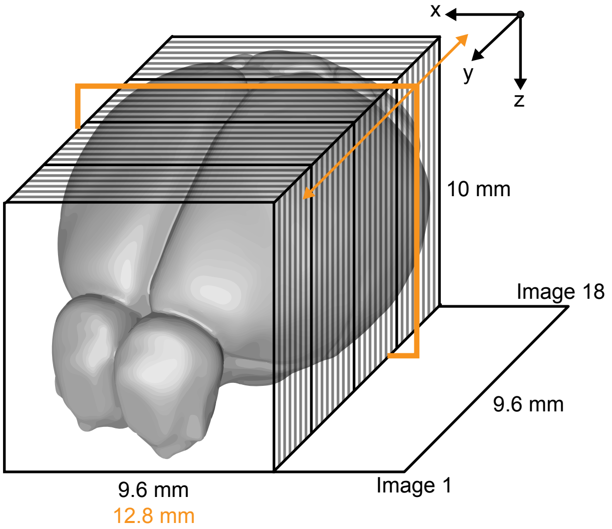


**Linear transducer**
128 piezo-elements - channels
Frequency: 15 MHz
8 mm focal (acoustic lens)
Voxel size: 100x300x100 $\mu m^3$
Field of view: 12.8x0.3x10 $mm^3$
Frame rate: 2-10 Hz
Cross-section μDoppler image

**2D-array transducer**
300-μm width between planes

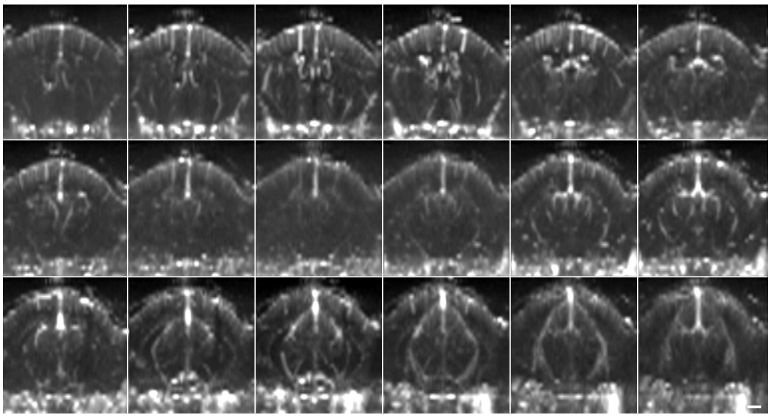

**Linear transducer**
300-μm step between planes

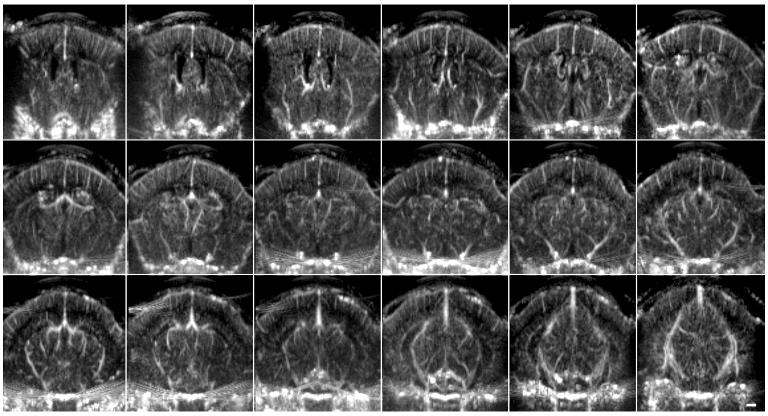

Table S1. List of brain regions in Figures 3C, 4C and 5C

| Region Number | Region Abbreviation | Brain region(s) | Name of the region | Volume of the region (mm3) |
|---|---|---|---|---|
| Cerebrum (CB) - Cortical Plate | | | | |
| 1 | GU | GU | Gustatory areas | 1.77 |
| 2 | SSp-bf | SSp-bf | Primary somatosensory area, barrel field | 6.29 |
| 3 | SSp-ll | SSp-ll | Primary somatosensory area, lower limb | 2.35 |
| 4 | SSp-tr | SSp-tr | Primary somatosensory area, trunk | 1.40 |
| 5 | SSp-m | SSp-m | Primary somatosensory area, mouth | 6.21 |
| 6 | SSp-n | SSp-n | Primary somatosensory area, nose | 3.02 |
| 7 | SSp-ul | SSp-ul | Primary somatosensory area, upper limb | 3.77 |
| 8 | SSp-un | SSp-un | Primary somatosensory area, unassigned | 1.26 |
| 9 | SSs | SSs | Supplemental somatosensory area | 9.03 |
| 10 | ACA | ACAd, ACAv | Anterior cingulate area | 5.51 |
| 11 | VISC | VISC | Visceral area | 2.35 |
| 12 | MOp | MOp | Primary motor area | 11.35 |
| 13 | MOs | MOs | Secondary motor area | 13.11 |
| 14 | AUDd | AUDd | Dorsal auditory area | 1.21 |
| 15 | AUDp | AUDp | Primary auditory area | 2.15 |
| 16 | AUDpo | AUDpo | Posterior auditory area | 0.61 |
| 17 | AUDv | AUDv | Ventral auditory area | 1.83 |
| 18 | VISa | VISa | Anterior visual area | 1.44 |
| 19 | VISrl | VISrl | Rostrolateral visual area | 1.02 |
| 20 | VISpor | VISpor | Postrhinal visual area | 1.25 |
| 21 | VISli | VISli | Laterointermediate visual area | 0.49 |
| 22 | VISal | VISal | Anterolateral visual area | 0.76 |
| 23 | VISam | VISam | Anteromedial visual area | 0.79 |
| 24 | VISpl | VISpl | Posterolateral visual area | 0.79 |
| 25 | VISpm | VISpm | Posteromedial visual area | 1.23 |
| 26 | VISl | VISl | Lateral visual area | 1.67 |
| 27 | VISp(pl) | VISp(pl) | Primary visual area, posterior lateral | 2.16 |
| 28 | VISp(pm) | VISp(pm) | Primary visual area, posterior medial | 1.04 |
| 29 | VISp(al) | VISp(al) | Primary visual area, anterior lateral | 1.34 |
| 30 | RSPagl | RSPagl | Restrosplenial area, lateral agranular part | 2.36 |
| 31 | RSPd | RSPd | Retrosplenial area, dorsal part | 3.80 |
| 32 | RSPv | RSPv | Retrosplenial area, ventral part | 4.33 |
| 33 | TEa | TEa | Temporal association areas | 3.11 |
| 34 | PERI | PERI | Perirhinal area | 0.79 |
| 35 | ECT | ECT | Ectorhinal area | 1.66 |

| | | | | |
|---|---|---|---|---|
| 36 | PIR | PIR | Piriform area | 11.57 |
| 37 | TR | TR | Postpiriform transition area | 1.40 |
| 38 | COA | COAa, COApl, COApm | Cortical amygdalar area | 3.27 |
| 39 | PAA | PAA | Piriform-amygdalar area | 1.19 |
| 40 | AI | AId, AIv | Agranular insular area | 5.47 |
| 41 | PL | PL | Prelimbic area | 2.40 |
| | Hippocampal Formation (HPF) | | | |
| 42 | CA1d CA2d CA3d | CA1d, CA2d, CA3d | CA1 CA2 CA3 subfields, dorsal part | 9.27 |
| 43 | CA1i | CA1i | CA1 subfield, intermediate | 2.05 |
| 44 | CA1v | CA1v | CA1 subfield, ventral | 2.07 |
| 45 | CA2i | CA2i | CA2 subfield, intermediate | 0.11 |
| 46 | CA2v | CA2v | CA2 subfield, ventral | 0.09 |
| 47 | CA3i | CA3i | CA3 subfield, intermediate | 1.80 |
| 48 | CA3v | CA3v | CA3 subfield, ventral | 1.56 |
| 49 | ENTl | ENTl | Entorhinal area, lateral part | 6.38 |
| 50 | ENTm | ENTm | Entorhinal area, medial part | 5.03 |
| 51 | PRE | PRE | Presubiculum | 0.92 |
| 52 | POST | POST | Postsubiculum | 1.08 |
| 53 | PAR | PAR | Parasubiculum | 0.93 |
| 54 | SUB | SUB | Subiculum | 2.10 |
| 55 | ProS | ProS | Prosubiculum | 1.30 |
| | Cerebrum (CB) - Cortical Subplate | | | |
| 56 | CLA | CLA | Claustrum | 0.55 |
| 57 | EP | Epd, EPv | Endopiriform nucleus | 2.79 |
| 58 | LA | LA | Lateral amygdalar nucleus | 0.84 |
| 59 | BLA | BLAa, BLAp, BLAv | Basolateral amygdalar nucleus | 1.90 |
| 60 | BMA | BMAa, BMAp | Basomedial amygdalar nucleus | 1.48 |
| 61 | PA | PA | Posterior amygdalar nucleus | 0.97 |
| | Striatum (STR) | | | |
| 62 | LSX | LSc, LSr, LSv | Lateral septal complex | 3.05 |
| 63 | CPadl | CPadl | Caudoputamen, anterior dorsal lateral | 1.00 |
| 64 | CPadm | CPadm | Caudoputamen, anterior dorsal medial | 1.80 |
| 65 | CPaml | CPaml | Caudoputamen, anterior medial lateral | 1.94 |
| 66 | CPamm | CPamm | Caudoputamen, anterior medial medial | 2.53 |
| 67 | CPavm | CPavm | Caudoputamen, anterior ventral medial | 1.12 |
| 68 | CPcdl | CPcdl | Caudoputamen, caudal dorsal lateral | 0.55 |
| 69 | CPcdm | CPcdm | Caudoputamen, caudal dorsal medial | 0.89 |
| 70 | CPcml | CPcml | Caudoputamen, caudal medial lateral | 2.17 |

| | | | | |
|---|---|---|---|---|
| 71 | CPcv | CPcv | Caudoputamen, caudal ventral | 0.75 |
| 72 | CPmdc | CPmdc | Caudoputamen, medial dorsal central | 2.25 |
| 73 | CPmdl | CPmdl | Caudoputamen, medial dorsal lateral | 0.36 |
| 74 | CPmdm | CPmdm | Caudoputamen, medial dorsal medial | 1.29 |
| 75 | CPmmc | CPmmc | Caudoputamen, medial medial central | 2.16 |
| 76 | CPmml | CPmml | Caudoputamen, medial medial lateral | 1.43 |
| 77 | CPmvc | CPmvc | Caudoputamen, medial ventral central | 2.03 |
| 78 | CPmvl | CPmvl | Caudoputamen, medial ventral lateral | 0.99 |
| 79 | ACB | ACB | Nucleus accumbens | 4.40 |
| 80 | OT | OT | Olfactory tubercle | 3.82 |
| 81 | sAMY | AAA, CEAc, CEAl, CEAm, IA, MEA | Striatum-like amygdalar nuclei | 4.02 |
| 82 | PAL | PAL | Pallidum | 1.13 |
| Thalamus (TH) | | | | |
| 83 | VAL | VAL | Ventral anterior-lateral complex of the thalamus | 0.82 |
| 84 | VM | VM | Ventral medial nucleus of the thalamus | 0.93 |
| 85 | VP | VPL, VPLpc, VPM, VPMpc | Ventral posterior complex of the thalamus | 2.83 |
| 86 | PP | PP | Peripeduncular nucleus | 0.06 |
| 87 | LGd | LGd-co, LGd-ip, LGd-sh | Dorsal part of the lateral geniculate complex | 0.63 |
| 88 | LP Eth | LP, Eth | Lateral posterior nucleus of the thalamus | 1.20 |
| 89 | PO | PO | Posterior complex of the thalamus | 1.25 |
| 90 | MD | MD | Mediodorsal nucleus of thalamus | 1.38 |
| 91 | RT | RT | Reticular nucleus of the thalamus | 1.45 |
| 92 | ATN | AD, Amd, Amv, AV, IAD, IAM, LD | Anterior group of the dorsal thalamus | 2.16 |
| 93 | MED | IMD, PR, SMT | Medial group of the dorsal thalamus | 0.63 |
| Hypothalamus (HY) | | | | |
| 94 | MEZ | AHN, PMd, LM, Mmd, Mml, Mmm, Mmme, Mmp, SUM, TMd, TMv, MPN, PVHd, PH, PMv, VMH | Hypothalamic medial zone | 3.84 |
| 95 | PVR | ADP, AVPV, AVP, DMH, MPO, MEPO, PS, PVp, PVpo, PD, SFO, SBPV, SCH, OV, | Periventricular region | 2.05 |
| 96 | ME | ME | Median eminence | 0.08 |
| 97 | LZ | LHA, LPO, PSTN, PeF, PST, RCH, STN, TU, | Hypothalamic lateral zone | 5.43 |
| 98 | PVZ | ASO, ARH, PVH, PVa, PVi, SO | Periventricular zone | 0.77 |
| Midbrain (MB) | | | | |
| 99 | SCd | SCd(a), SCd(p) | Superior colliculus, deep layers | 1.52 |

| | | | | |
|---|---|---|---|---|
| 100 | SCi | SCi(al), SCi(am) | Superior colliculus, intermediate layers | 3.58 |
| 101 | SCi(pl) | SCi(pl) | Superior colliculus, intermediate layers, posterior lateral | |
| 102 | SCs(a) | SCs(am), SCs(al) | Superior colliculus, superficial layers, anterior | 0.96 |
| 103 | SCs(pl) | SCs(pl) | Superior colliculus, superficial layers, posterior lateral | 0.59 |
| 104 | SCs(pm) | SCs(pm) | Superior colliculus, superficial layers, posterior medial | 0.61 |
| 105 | ICc | ICc | Inferior colliculus, central nucleus | 1.13 |
| 106 | ICd | ICd | Inferior colliculus, dorsal nucleus | 1.32 |
| 107 | ICe | ICe | Inferior colliculus, external nucleus | 2.00 |
| 108 | MRN | MRNal, MRNam, MRNpl, MRNpm | Midbrain reticular nucleus | 5.14 |
| 109 | CUN | CUN | Cuneiform nucleus | 0.55 |
| 110 | SN | SNr, SNc | Substantia nigra | 1.55 |
| 111 | PAG | PAGa, PAGmd, PAGml, PAGmv, PAGpd, PAGpl, PAGpv | Periaqueductal gray | 4.30 |
| 112 | APN | APN | Anterior pretectal nucleus | 1.28 |
| 113 | RN | RN | Red nucleus | 0.79 |
| 114 | PPN | PPN | Pedunculopontine nucleus | 0.89 |
| Hindbrain (HB) | | | | |
| 115 | NLL | NLL | Nucleus of the lateral lemniscus | 0.72 |
| 116 | PSV | PSV | Principal sensory nucleus of the trigeminal | 1.10 |
| 117 | PB | PB | Parabrachial nucleus | 0.95 |
| 118 | SOC | SOCl, SOCm | Superior olivary complex | 0.53 |
| 119 | PRNc | PRNc | Pontine reticular nucleus, caudal part | 2.35 |
| 120 | PCG | PCG | Pontine central gray | 0.54 |
| 121 | PG | PG | Pontine gray | 0.96 |
| 122 | TRN | TRN | Tegmental reticular nucleus | 0.69 |
| 123 | CS | CS | Superior central nucleus raphe | 0.59 |
| 124 | PRNr | PRNr | Pontine reticular nucleus | 2.36 |
| 125 | VII | VII | Facial motor nucleus | 0.92 |
| 126 | GRN | GRN | Gigantocellular reticular nucleus | 2.61 |
| 127 | IRN | IRN | Intermediate reticular nucleus | 2.78 |
| 128 | MARN | MARN | Magnocellular reticular nucleus | 0.53 |
| 129 | PARN | PARN | Parvicellular reticular nucleus | 2.25 |
| Cerebellum (CBL) | | | | |
| 130 | CENT | CENT2, CENT3 | Central lobule | 4.05 |
| 131 | SIM | SIM | Simple lobule | 5.69 |
| 132 | CUL | CUL4,5 | Lobules IV-V | 6.71 |

| | | | | |
|---|---|---|---|---|
| 133 | PFL | PFL | Paraflocculus | 5.72 |
| 134 | FL | FL | Flocculus | 1.34 |

Table S2. List of brain regions for RSFC in Figure S3

| Region Number | Region Abbreviation | Brain region(s) | Name of the region | Volume of the region (mm3) |
|---|---|---|---|---|
| Cerebrum (CB) - Cortical Plate | | | | |
| 1 | TR | TR | Postpiriform transition area | 1.40 |
| 2 | TEa | TEa | Temporal association areas | 3.11 |
| 3 | AUDv | AUDv | Ventral auditory area | 1.83 |
| 4 | SSs | SSs | Supplemental somatosensory area | 9.03 |
| 5 | SSp-n | SSp-n | Primary somatosensory area, nose | 3.02 |
| 6 | SSp-m | SSp-m | Primary somatosensory area, mouth | 6.21 |
| 7 | SSp-bf | SSp-bf | Primary somatosensory area, barrel field | 6.29 |
| 8 | MOp | MOp | Primary motor area | 11.35 |
| 9 | VISli | VISli | Laterointermediate visual area | 0.49 |
| 10 | PAA | PAA | Piriform-amygdalar area | 1.19 |
| 11 | COA | COAa, COApl, COApm | Cortical amygdalar area | 3.27 |
| 12 | VISrl | VISrl | Rostrolateral visual area | 1.02 |
| 13 | SSp-un | SSp-un | Primary somatosensory area, unassigned | 1.26 |
| 14 | SSp-ul | SSp-ul | Primary somatosensory area, upper limb | 3.77 |
| 15 | MOs | MOs | Secondary motor area | 13.11 |
| 16 | AUDd | AUDd | Dorsal auditory area | 1.21 |
| 17 | VISp(al) | VISp(al) | Primary visual area, anterior lateral | 1.34 |
| 18 | VISl | VISl | Lateral visual area | 1.67 |
| 19 | VISp(pl) | VISp(pl) | Primary visual area, posterior lateral | 2.16 |
| 20 | VISp(pm) | VISp(pm) | Primary visual area, posterior medial | 1.04 |
| 21 | VISpm | VISpm | Posteromedial visual area | 1.23 |
| 22 | RSPagl | RSPagl | Retrosplenial area, lateral agranular part | 2.36 |
| 23 | RSPd | RSPd | Retrosplenial area, dorsal part | 3.80 |
| 24 | SSp-ll | SSp-ll | Primary somatosensory area, lower limb | 2.35 |
| 25 | SSp-tr | SSp-tr | Primary somatosensory area, trunk | 1.40 |
| 26 | VISam | VISam | Anteromedial visual area | 0.79 |
| 27 | VISa | VISa | Anterior visual area | 1.44 |
| 28 | ACA | ACAd, ACAv | Anterior cingulate area | 5.51 |
| 29 | RSPv | RSPv | Retrosplenial area, ventral part | 4.33 |
| 30 | PL | PL | Prelimbic area | 2.40 |
| 31 | VISal | VISal | Anterolateral visual area | 0.76 |
| 32 | AUDpo | AUDpo | Posterior auditory area | 0.61 |
| 33 | AUDp | AUDp | Primary auditory area | 2.15 |
| 34 | VISpl | VISpl | Posterolateral visual area | 0.79 |
| 35 | VISpor | VISpor | Postrhinal visual area | 1.25 |
| 36 | PIR | PIR | Piriform area | 11.57 |
| 37 | PERI | PERI | Perirhinal area | 0.79 |
| 38 | VISC | VISC | Visceral area | 2.35 |
| 39 | GU | GU | Gustatory areas | 1.77 |
| 40 | AI | AId, AIv | Agranular insular area | 5.47 |
| 41 | ECT | ECT | Ectorhinal area | 1.66 |
| Thalamus (TH) | | | | |
| 42 | RT | RT | Reticular nucleus of the thalamus | 1.45 |
| 43 | VM | VM | Ventral medial nucleus of the thalamus | 0.93 |
| 44 | MD | MD | Mediodorsal nucleus of thalamus | 1.38 |
| 45 | MED | IMD, PR, SMT | Medial group of the dorsal thalamus | 0.63 |
| 46 | PP | PP | Peripeduncular nucleus | 0.06 |
| 47 | LGd | LGd-co, LGd-ip, LGd-sh | Dorsal part of the lateral geniculate complex | 0.63 |
| 48 | ATN | AD, Amd, Amv, AV, IAD, IAM, LD | Anterior group of the dorsal thalamus | 2.16 |
| 49 | VAL | VAL | Ventral anterior-lateral complex of the thalamus | 0.82 |
| 50 | VP | VPL, VPLpc, VPM, VPMpc | Ventral posterior complex of the thalamus | 2.83 |
| 51 | PO | PO | Posterior complex of the thalamus | 1.25 |
| 52 | LP Eth | LP, Eth | Lateral posterior nucleus of the thalamus | 1.20 |
| Hindbrain (HB) | | | | |
| 53 | VII | VII | Facial motor nucleus | 0.92 |
| 54 | SOC | SOCl, SOCm | Superior olivary complex | 0.53 |
| 55 | PG | PG | Pontine gray | 0.96 |
| 56 | TRN | TRN | Tegmental reticular nucleus | 0.69 |
| 57 | MARN | MARN | Magnocellular reticular nucleus | 0.53 |
| 58 | NLL | NLL | Nucleus of the lateral lemniscus | 0.72 |

| | | | | |
|---|---|---|---|---|
| 59 | PRNc | PRNc | Pontine reticular nucleus, caudal part | 2.35 |
| 60 | PRNr | PRNr | Pontine reticular nucleus | 2.36 |
| 61 | GRN | GRN | Gigantocellular reticular nucleus | 2.61 |
| 62 | PARN | PARN | Parvicellular reticular nucleus | 2.25 |
| 63 | IRN | IRN | Intermediate reticular nucleus | 2.78 |
| 64 | PCG | PCG | Pontine central gray | 0.54 |
| 65 | PB | PB | Parabrachial nucleus | 0.95 |
| 66 | PSV | PSV | Principal sensory nucleus of the trigeminal | 1.10 |
| 67 | CS | CS | Superior central nucleus raphe | 0.59 |
| Cerebellum (CBL) | | | | |
| 68 | FL | FL | Flocculus | 1.34 |
| 69 | PFL | PFL | Paraflocculus | 5.72 |
| 70 | CUL | CUL4,5 | Lobules IV-V | 6.71 |
| 71 | CENT | CENT2 CENT3 | Central lobule | 4.05 |
| 72 | SIM | SIM | Simple lobule | 5.69 |
| Midbrain (MB) | | | | |
| 73 | SN | SNr, SNc | Substantia nigra | 1.55 |
| 74 | PAG | PAGa, PAGmd, PAGml, PAGmv, PAGpd, PAGpl, PAGpv | Periaqueductal gray | 4.30 |
| 75 | MRN | MRNal, MRNam, MRNpl, MRNpm | Midbrain reticular nucleus | 5.14 |
| 76 | PPN | PPN | Pedunculopontine nucleus | 0.89 |
| 77 | ICe | ICe | Inferior colliculus, external nucleus | 2.00 |
| 78 | ICd | ICd | Inferior colliculus, dorsal nucleus | 1.32 |
| 79 | SCi | SCi(al), SCi(am) | Superior colliculus, intermediate layers | 3.58 |
| 80 | SCi(pl) | SCi(pl) | Superior colliculus, intermediate layers, posterior lateral | 0,59 |
| 81 | SCs(pm) | SCs(pm) | Superior colliculus, superficial layers, posterior medial | 0.61 |
| 82 | SCs(a) | SCs(am), SCs(al) | Superior colliculus, superficial layers, anterior | 0.96 |
| 83 | SCd | SCd(a), SCd(p) | Superior colliculus, deep layers | 1.52 |
| 84 | ICc | ICc | Inferior colliculus, central nucleus | 1.13 |
| 85 | CUN | CUN | Cuneiform nucleus | 0.55 |
| 86 | APN | APN | Anterior pretectal nucleus | 1.28 |
| 87 | RN | RN | Red nucleus | 0.79 |
| 88 | SCs(pl) | SCs(pl) | Superior colliculus, superficial layers, posterior lateral | 0.59 |
| Hypothalamus (HY) | | | | |
| 89 | PVZ | ASO, ARH, PVH, PVa, PVi, SO | Periventricular zone | 0.77 |
| 90 | ME | ME | Median eminence | |
| 91 | PVR | ADP, AVPV, AVP, DMH, MPO, MEPO, PS, PVp, PVpo, PD, SFO, SBPV, SCH, OV, VLPO, VMPO | Periventricular region | 2.05 |
| 92 | LZ | LHA, LPO, PSTN, PeF, PST, RCH, STN, TU, ZI | Hypothalamic lateral zone | 5.43 |
| 93 | MEZ | AHN, PMd, LM, Mmd, Mml, Mmm, Mmme, Mmp, SUM, TMd, TMv, MPN, PVHd, PH, PMv, | Hypothalamic medial zone | 3.84 |
| Striatum (STR) | | | | |
| 94 | LSX | LSc, LSr, LSv | Lateral septal complex | 3.05 |
| 95 | CPmdm | CPmdm | Caudoputamen, medial dorsal medial | 1.29 |
| 96 | CPadl | CPadl | Caudoputamen, anterior dorsal lateral | 1.00 |
| 97 | CPadm | CPadm | Caudoputamen, anterior dorsal medial | 1.80 |
| 98 | OT | OT | Olfactory tubercle | 3.82 |
| 99 | CPamm | CPamm | Caudoputamen, anterior medial medial | 2.53 |
| 100 | CPcdm | CPcdm | Caudoputamen, caudal dorsal medial | 0.89 |
| 101 | CPcdl | CPcdl | Caudoputamen, caudal dorsal lateral | 0.55 |
| 102 | CPcml | CPcml | Caudoputamen, caudal medial lateral | 2.17 |
| 103 | CPmdc | CPmdc | Caudoputamen, medial dorsal central | 2.25 |
| 104 | CPaml | CPaml | Caudoputamen, anterior medial lateral | 1.94 |
| 105 | CPmdl | CPcml | Caudoputamen, caudal medial lateral | 2.17 |
| 106 | PAL | PAL | Pallidum | 1.13 |
| 107 | CPcv | CPcv | Caudoputamen, caudal ventral | 0.75 |
| 108 | CPmml | CPmml | Caudoputamen, medial medial lateral | 1.43 |
| 109 | CPmmc | CPmmc | Caudoputamen, medial medial central | 2.16 |
| 110 | CPavm | CPavm | Caudoputamen, anterior ventral medial | 1.12 |

| | | | | |
|---|---|---|---|---|
| 111 | sAMY | AAA, CEAc, CEAl, CEAm, IA, | Striatum-like amygdalar nuclei | 4.02 |
| 112 | CPmvl | CPmvl | Caudoputamen, medial ventral lateral | 0.99 |
| 113 | CPmvc | CPmvc | Caudoputamen, medial ventral central | 2.03 |
| 114 | ACB | ACB | Nucleus accumbens | 4.40 |
| Cerebrum (CB) - Cortical Subplate + Hippocampal Formation | | | | |
| 115 | EP | EPd, Epv | Endopiriform nucleus | 2.79 |
| 116 | LA | LA | Lateral amygdalar nucleus | 0.84 |
| 117 | CA1i | CA1i | CA1 subfield, intermediate | 2.05 |
| 118 | BLA | BLAa, BLAp, BLAv | Basolateral amygdalar nucleus | 1.90 |
| 119 | CA3i | CA3i | CA3 subfield, intermediate | 1.80 |
| 120 | CA2i | CA2i | CA2 subfield, intermediate | 0.11 |
| 121 | CLA | CLA | Claustrum | 0.55 |
| 122 | BMA | BMAa, BMAp | Basomedial amygdalar nucleus | 1.48 |
| 123 | CA1v | CA1v | CA1 subfield, ventral | 2.07 |
| 124 | CA3v | CA3v | CA3 subfield, ventral | 1.56 |
| 125 | PA | PA | Posterior amygdalar nucleus | 0.97 |
| 126 | ENTl | ENTl | Entorhinal area, lateral part | 6.38 |
| 127 | ENTm | ENTm | Entorhinal area, medial part | 5.03 |
| 128 | ProS | ProS | Prosubiculum | 1.30 |
| 129 | CA2v | CA2v | CA2 subfield, ventral | 0.09 |
| 130 | POST | POST | Postsubiculum | 1.08 |
| 131 | PAR | PAR | Parasubiculum | 0.93 |
| 132 | PRE | PRE | Presubiculum | 0.92 |
| 133 | SUB | SUB | Subiculum | 2.10 |
| 134 | CA1d CA2d CA3d | CA1d, CA2d, CA3d | CA1 CA2 CA3 subfields, dorsal part | 9.27 |
