## Supplementary code 1 - Registration for "A platform for brain-wide functional ultrasound imaging and analysis of circuit dynamics in behaving mice": exampleDisplay.m

```
%%% DISPLAY

load('dataAtlas');
load('dataExample');
load('rotationMatrixExample')
 
dopplerReg=registerData(doppler,mvr);
 
figure(1);
imagesc(squeeze(dopplerReg(140,:,:)))
colormap('gray')
caxis([0.4 0.8])
addLines(LinReg.Cor,140);

figure(2);
imagesc(squeeze(dopplerReg(:,40,:)))
colormap('gray')
caxis([0.4 0.8])
addLines(LinReg.Tra,40);
```
