## Supplementary code 1 - Registration for "A platform for brain-wide functional ultrasound imaging and analysis of circuit dynamics in behaving mice": exampleRegistration.m

```
%%% REGISTRATION

load dataAtlas;
load dataExample;
MatlasReg=mapscan(AtlasReg,sa.rgb,'index');
MatlasHis=mapscan(AtlasHis,gray(64),'auto');
Mdoppler=mapscan(doppler,gray(64),'fix');
Mdoppler.caxis=[0.4 0.8];

register3D(Mdoppler,MatlasHis,LinReg);

 
```
