## Supplementary code 1 - Registration for "A platform for brain-wide functional ultrasound imaging and analysis of circuit dynamics in behaving mice": exampleSegmentation.m

```
%%% SEGMENTATION

load('dataAtlas');
load('dataExample');
load('rotationMatrixExample')

% compute the mask
dopplerReg=registerData(doppler,mvr);
nreg=acr2num(sa,'SSp-bf');
maskRegion=dopplerReg.*(AtlasReg==nreg);

% display the result
figure;
imagesc(squeeze(maskRegion(140,:,:)))
colormap('gray')
caxis([0.4 0.8])

% display the total intensity in the region
fprintf('total intensity in SSp-bf %f AU\n', sum(maskRegion(:)));
```
