## Supplementary code 1 - Registration for "A platform for brain-wide functional ultrasound imaging and analysis of circuit dynamics in behaving mice": README.txt

Note : the following sections use the same example data ("dataExample.mat"), atlas ("dataAtlas.mat") and rotation matrix ("rotationMatrixExample.mat")


REGISTRATION

Script to run : exampleRegistration.m

Tested Matlab version : R2017b

Description : provides a grapical user interface to perform the 3D data registration (commands below)
      * "caxis" : set the color limits (example : 0.4 0.8)
      * "colormap" : select a colormap (example : gray)
      * left-clic dragging : translate one of the microdoppler images
      * right-clic dragging : rotate one of the microdoppler images
      * "apply": after performing a translation/rotation, use this button to rotate the full 3D data in the other planes
      * "save": save the computed parameters in the file rotateMatrix.mat
      * sliders beside view : navigation in the data along the associated axis

Important notes : 1. the user should use the "apply" button every time a translation/rotation is performed on a view
		  2. better registration is achieved when adjusting the data to the atlas at different spatial locations
		  3. it may happen that some translation/rotation is not taken into account while clicking on "apply". To solve this issue,
			please use the sliders to move to another location in the data.


DISPLAY

Script to run : exampleDisplay.m

Tested Matlab version : R2017b

Description : provides 2 displays of the registered data with atlas boundaries on overlay


SEGMENTATION

Script to run : exampleSegmentation.m

Tested Matlab version : R2017b

Description : 1. computes a region-specific mask on the data (in the example the somatosensory cortex SSp-bf)
	      2. displays the data with the mask
	      3. computes the average intensity in the region
