## Supplementary code 2 - Retinotopic mapping for "A platform for brain-wide functional ultrasound imaging and analysis of circuit dynamics in behaving mice": README.txt

RETINOTOPIC MAPPING

Script to run : exampleRetinotopicMapping.m 

Tested Matlab version : R2017b

Description : 1. load the example data "dataRetinotopicMapping.mat"
	      2. computes the delay of the maximum response
	      3. display the microdoppler image and the resulting delay map
 
