## Supplementary code 2 - Retinotopic mapping for "A platform for brain-wide functional ultrasound imaging and analysis of circuit dynamics in behaving mice": retinotopy.m

```
function r=retinotopie(Im)

[nz,nxy,nt]=size(Im);
r=zeros(nt,nxy);
a=hanning(7);
for iz=1:nz
    for ixy=1:nxy
       s=squeeze(Im(iz,ixy,:));
       x=xcorr(s,a);
       x=interp1(double(x),[0:1/16:length(x)],'PCHIP');
       [~,im]=max(x);
       r(iz,ixy)=im/16-nt;
    end
end
end
```
