## Supplementary code 3 - Resting state functional connectivity for "A platform for brain-wide functional ultrasound imaging and analysis of circuit dynamics in behaving mice": correlation.m

```
% basic correlation 
% b(x,y,t)  3D matrix 
% a(t) signal to correlate with

function c=correlation(b,a)
[nz,nx,nt]=size(b); 

a=a-mean(a); 
a=a./sqrt(sum(a.^2));
 
c=zeros(nz,nx);
 
 for iz=1:nz
     for ix=1:nx
          s=squeeze(b(iz,ix,:));
          s=s-mean(s);
          s=s/sqrt(sum(s.^2));
          c(iz,ix)=sum(s.*a);
     end
 end

 
```
