## Supplementary code 3 - Resting state functional connectivity for "A platform for brain-wide functional ultrasound imaging and analysis of circuit dynamics in behaving mice": exampleRestingState.m

```
%%% RESTING STATE FUNCTIONAL CONNECTIVITY

load('dataRestingState');

% apply the filters
Irf=filterNoise(Ir,15);
Irf=filterOutlier(Irf,1.8);
Fs=1/(15*0.7);
[B,A]=butter(2,0.002/Fs*2,'high');
Irf=filter(B,A,Irf,[],3);
Irf=filterAverage(Irf);   
Irf=normalize(Irf);

% compute the correlation with a seed at location (6, 43) and display
seed=squeeze(Irf(6,43,:));
c=correlation(Irf,seed); 
imagesc(c);
caxis([-0.7 0.7])
colormap jet;
title('seed 6 43');
```
