## Supplementary code 3 - Resting state functional connectivity for "A platform for brain-wide functional ultrasound imaging and analysis of circuit dynamics in behaving mice": filterAverage.m

```
function Imf=filterAverage(Im)

[nz,nxy,nt]=size(Im);

Imf=zeros(nz,nxy,nt);

st=squeeze(mean(mean(Im)));
st=st-mean(st);
st=st./sqrt(sum(st.^2));

for iz=1:nz
    for ixy=1:nxy
      tmp=squeeze(Im(iz,ixy,:));
      tmp=tmp-st*sum(st.*tmp);  % substraction of the average signal
      Imf(iz,ixy,:)=tmp;
    end
end

end
```
