## Supplementary code 3 - Resting state functional connectivity for "A platform for brain-wide functional ultrasound imaging and analysis of circuit dynamics in behaving mice": filterNoise.m

```
function Im=filterNoise(Ie,n)

[nz,nx,nt]=size(Ie);

ntr=floor(nt/n)-2;
n2=floor(n/3);
Im=zeros(nz,nx,ntr);
ss=squeeze(mean(mean(Ie)));

for i=1:ntr 
    tmp=ss((1:n)+(i-1)*n);
    [~,ind]=sort(tmp);   
    Im(:,:,i)=mean(Ie(:,:,ind(1:n2)+(i-1)*n),3); 
end

end
```
