## Supplementary code 3 - Resting state functional connectivity for "A platform for brain-wide functional ultrasound imaging and analysis of circuit dynamics in behaving mice": normalize.m

```
function Im=normalize(Ix)

[nz,nxy,nt]=size(Ix);
Im=zeros(nz,nxy,nt);

for iz=1:nz
    for ix=1:nxy
        val=squeeze(Ix(iz,ix,:));
        val=(val-mean(val))./(sqrt(sum(val.^2))+1e-10);
        Im(iz,ix,:)=val;
    end
end

end
```
