## Supplementary code 3 - Resting state functional connectivity for "A platform for brain-wide functional ultrasound imaging and analysis of circuit dynamics in behaving mice": README.txt

RESTING STATE FUNCTIONAL CONNECTIVITY

Script to run : exampleRestingState.m

Tested Matlab version : R2017b

Description : 1. load the example data "dataRestingState.mat"
	      2. apply the sequence of filters
	      3. compute and display the map of the correlation with the seed at coordinates (6, 43)
